# Longitudinal preclinical imaging characterization of drug delivery potential after radiotherapy in the healthy and leukemic bone marrow vascular microenvironment

**DOI:** 10.1101/2021.02.23.432514

**Authors:** Jamison Brooks, Darren Zuro, Joo Y. Song, Srideshikan Sargur Madabushi, James F Sanchez, Marcin Kortylewski, Bihong T. Chen, Kalpna Gupta, Guy Storme, Jerry Froelich, Susanta K Hui

## Abstract

**Objectives:** Radiotherapy improves blood perfusion and cellular chemotherapy uptake in mice with acute lymphoblastic leukemia (ALL). However, its ability to influence drug delivery and permeation through the bone marrow vasculature (BMV) is unknown, due in part to a lack of methodology. This study developed longitudinal quantitative multiphoton (L-QMPM) imaging and used it to characterize drug delivery potential and the BMV before and after radiotherapy in mice bearing leukemia.

**Methods:** We developed a longitudinal window implant for L-QMPM imaging of the calvarium BMV before, 2 days after, and 5 days after radiotherapy. Live time-lapsed images of a fluorescent drug surrogate were used to obtain measurements including tissue wash-in slope (WIS_tissue_) to measure drug delivery potential. We performed L-QMPM imaging using 2 Gy and 10 Gy total body irradiation (TBI) on C57/B6 (WT) mice, mice bearing ALL, and acute myeloid leukemia (AML).

**Results:** Implants had no effects on calvarium dose, and parameters for WT untreated mice were stable during imaging. We observed increased angiogenesis, decreased single-vessel blood flow, and decreased WIS_tissue_ with the onset of AML and ALL. 2Gy and 10Gy TBI increased WIS_tissue_ 2 days after radiotherapy in all 3 groups of mice and increased single-vessel blood flow in mice bearing ALL and AML. Significant increases in WIS_tissue_ were observed 2 days after 2Gy TBI compared to 5 days. Morphological and functional alterations in the BMV were sustained for a significantly longer time period after 10Gy TBI (5 days post-treatment) compared to 2Gy TBI (2 days post-treatment).

**Conclusion:** L-QMPM provides stable functional assessments of the BMV. TBI increases the drug delivery potential of the leukemic BMV 2-5 days post-treatment, likely through improved blood perfusion and drug exchange from the BMV to the extravascular tissue. Our data show that neo-adjuvant 2Gy and 10Gy TBI condition the BMV for increased drug delivery.

## Introduction

It has been shown that leukemia induced bone marrow vasculature (BMV) remodeling promotes chemoresistance in mice bearing acute myeloid leukemia (AML) (1,2). Angiogenesis is commonly observed in patients with leukemia (3), and murine studies using human AML have reported vascular alterations that result in poor drug tissue perfusion, similarly to solid tumor models (4). Despite such observations, anti-angiogneic therapeutic approaches in combination with standard chemotherapy have not been successful in improving patient outcomes for leukemia (5,6). This may in part be due to the complex response of the vascular system to treatment as well as a lack of methods to assess the function of the BMV (7).

One technique to assess the function of the BMV is dynamic contrast-enhanced (DCE) imaging with modalities such as magnetic resonance imaging (MRI) or computed tomography (CT). DCE imaging acquires time-lapsed images of intravenously injected contrast agents to quantify changes in their delivery to the tissue (8). However, DCE imaging is unable to directly observe the underlying vascular characteristics influencing changes in contrast delivery(9). DCE imaging using a small molecular weight contrast agent (938 Da) has revealed increased peak height and wash-in slope in the bone marrow of patients with AML compared to healthy volunteers (10). Increased blood perfusion and blood volume have been proposed as reasons for changes in peak height and wash-in slope, respectively. Other studies have noticed increased clearance rates of compounds from the bone marrow of patients with acute leukemias (11); however, the results of these experiments are very different from previously mentioned preclinical observations of poor chemotherapy delivery with the onset of leukemia. Such apparent differences between preclinical and clinical observations demonstrate the need for preclinical methodology capable of performing clinically relevant contrast-based time-lapsed imaging, while directly observing BMV alterations.

Multiphoton microscopy is one imaging technique capable of both directly observing the BMV and performing contrast-based time-lapsed imaging. Fluorescent dextran conjugates of a variety of molecular weights are commonly used for multiphoton microscopy as a contrast agent. d leakage of dextran with molecular weights up to 150 kDa have been observed in the bone marrow (1,12) due to the presence of highly permeable sinusoidal vessels (13). In comparison to 150 kDa sized dextran, 4kDa sized dextran easily permeates the BMV and is similar in molecular weight to typical chemotherapies used in the treatment of leukemia such as daunorubicin (563.98 Da) or cytarabine (243.22 Da) (14). It is also similar in size to previously mentioned contrast agents used in DCE imaging. Compared to chemotherapy, 4kDa dextran has a minimal effect on the bone marrow microenvironment (15), making it ideal for longitudinal studies as a drug surrogate and useful for comparison with DCE imaging.

A recent study using multiphoton microscopy found that low dose (2-4 Gy) radiotherapy increases single-vessel blood flow in mice bearing acute lymphoblastic leukemia (ALL) (12). Increases in perfusion coincided with increased cellular uptake of chemotherapy, as well as improved survival outcome for mice treated with chemotherapy after neo-adjuvant low-dose radiotherapy. The study suggests that improved BMV blood flow may improve drug delivery to the bone marrow. However, the correlation between improved BMV blood flow and improved cellular uptake of chemotherapy does not assure increased drug delivery potential after radiotherapy. Additional factors influence cellular uptake of therapeutics, such as tissue cellularity and the affinity of cells to uptake chemotherapy (16). Furthermore, measurements of cellular drug uptake are limited to single time-point measurements, and do not have the ability to observe longitudinal changes required for sustained drug delivery over a period of time. Therefore, longitudinal measurements of drug delivery potential would be beneficial to assess the effectiveness of radiotherapy in conditioning the BMV for chemotherapy delivery.

We have developed longitudinal quantitative multiphoton microscopy (L-QMPM) to, 1) directly assess the drug delivery potential of the BMV after radiotherapy, 2) quantify potential time-lapsed imaging-based disease biomarkers, and 3) better understand the relationship between underlying vascular alterations and measurements obtained from DCE imaging. We quantified drug delivery potential primarily as the extravascular tissue wash-in slope (WIS_tissue_) calculated usingtime-lapsed imaging of 4kDa dextran. We measured several additional time-lapsed imaging and microvascular parameters, including the kinetic transfer rate (K_trans_) and single-vessel blood flow to understand how the BMV influences changes in DCE imaging parameters and WIS_tissue_. We performed L-QMPM imaging in the calvarium of mice bearing ALL or AML, two common types of leukemia, as well as wild-type (WT) controls, to assess the response of the healthy and leukemic BMV to radiotherapy. We hypothesized that radiotherapy would increase WIS_tissue_ in the calvarium bone marrow of mice bearing leukemia after both low (2 Gy) and high (10 Gy) dose total body irradiation (TBI).

## Methods

### Mice and cell lines

We performed all procedures for animal experimentation according to City of Hope guidelines and approved by the Institutional Animal Care and Use Committee. For all imaging studies, we used male and female C57BL/6 WT mice (Strain 556, Charles River, Wilmington MA USA) at 7-11 weeks. Mice were injected with 1×10^6^ green fluorescent protein (GFP)^+^ BCR-ABL (p190Kd) expressing B-cell ALL cells (17) or 1×10^6^ GFP^+^ *Cbfb-MYH11/Mpl*^+^ AML cells (18). The percentage of leukemia in peripheral blood (PB) and in bone samples was measured by flow cytometry as described previously (12). Complete details of the mice and leukemia injections can be found in Appendix 1:A

### Cranial Window Surgery

Briefly, we anesthetized mice using 1.3% isoflurane, and made an incision to expose the calvarium. A custom head plate made of carbon fiber or titanium for sterotactic viewing of the calvarium and a round glass coverslip were adhered to the calvarium using a combination of adhesives. Complete details of the surgery can be found in Appendix 1:B.

### L-QMPM image acquisition and analysis

For L-QMPM imaging, we anesthetized mice using 1.3% isoflurane. The cranial window head plate attached to the mouse was inserted into a stereotactic heated stage for imaging using a Prairie Ultima multiphoton microscope (Bruker Corporation, Billica MA USA). We performed maging on the frontal bone region of the calvarium, near the sagittal suture for all mice. For time-lapsed contrast imaging, we inserted a catheter into the tail vein of the mice (19) and injected 4 kDa TRITC dextran (TdB Consultancy, Uppsala Sweden) during time-lapsed image acquisition. A second injection of 2 MDa FITC dextran (TdB Consultancy, Uppsala Sweden) or Qtracker™ 655 Vascular Label (Invitrogen, Carlsbad CA USA) was performed to observe the vasculature morphology and identify vascular tissue. We used three separate z-stack images to measure vessel density and mean vessel diameter for each mouse using Fiji/Imagej (20). Measurements of corrected single-vessel blood flow were acquired as described previously (4). Tiled z-stack images were stitched together using a Fiji/Imagej plugin (21). A complete description of image acquisition can be found in Appendix 1:C.

For identification of extravascular, vascular, and whole tissue regions of interest (ROI), we calculated a series of automatic thresholds from fluorescent blood pool agents and time-lapsed 4 kDa dextran images using Otsu’s method (22) in Fiji/ImageJ. Time-lapsed dextran accumulation in the vascular and extravascular tissue ROIs were analyzed using a custom Matlab® (R2018a 9.41.0.81364, MathWorks Natick, MA) script to obtain K_trans_, the fractional extracellular extravascular space (ν_ec_), and the backflux rate (K_ep_) using a 2-compartment model (23) as described prevously (12). The additional parameters of dextran accumulation that we quantified included WIS_tissue_, wash-in slope for the whole tissue ROI (WIS_whole_), peak height for the extravascular tissue ROI (PH_tissue_), peak height for the whole tissue ROI (PH_whole_), and peak height for the vascular tissue ROI (PH_blood_). WIS_whole_ and PH_whole_ were quantified as measurements similar to DCE imaging parameters to identify how signal contributions from the vascular and extravascular tissue influence DCE imaging(9).

We obtained WIS_tissue_ and WIS_whole_ by dividing the largest positive change in fluorescent intensity between each frame by the time interval between frames for the ROIs of the extravascular and whole tissues, respectively. PH_blood_, PH_tissue_, and PH_whole_ were calculated by first subtracting the pre-injection background fluorescence from time-lapsed images, then measuring the dextran fluorescent intensity value of the frame with the highest fluorescent intensity for vascular, extravascular, and whole tissue ROIs, respectively (8) (Figure 1A-B). A complete description of image segmentation, compartmental modeling, and descriptive curve analysis is found in the Appendix 1:D.

**Figure 1:**
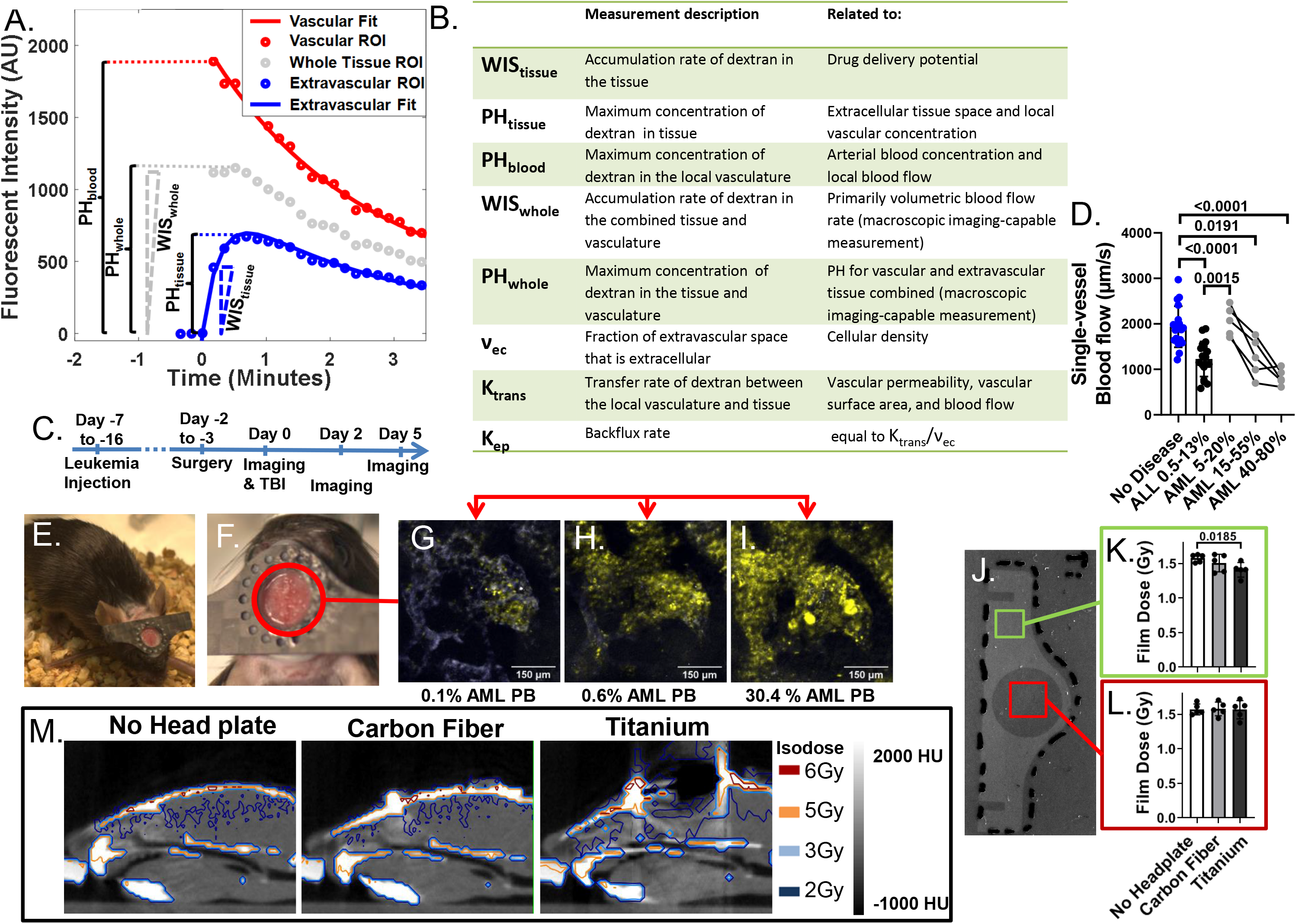
Time-lapsed imaging parameters and validation of longitudinal cranial window head plates. Fluorescent intensity values from extravascular tissue, vascular tissue, and whole tissue ROIs from time-lapsed L-QMPM images. Solid lines represent fitted functions for the vascular and extravascular tissue compartments used to obtain K_trans_,K_ep_, and ν_ec_. WIS_tissue_ and WIS_whole_ are depicted as the maximum positive slope between image frames for the extravascular tissue ROI and whole tissue ROI, respectively. A summary of imaging parameters used in this study. (C) The L-QMPM imaging schema. (D) Single-vessel blood flow values for WT mice and mice with varying percentages of leukemia in PB. Mice bearing ALL were imaged 7-8 days after ALL injection and mice bearing AML were imaged longitudinally 9, 11, and 14 days after AML injection. (E-F) Photos of mice with longitudinal cranial windows. L-QMPM Images of GFP+ AML cell growth (Green) in the calvarium at (G) 4, (H) 6, and (I) 10 days post AML injection shown with corresponding peripheral blood measurements. (J) A picture of dosimetric film after x-ray exposure while directly underneath a titanium head plate. Film dose measurements from a lack of head plate, carbon fiber head plates, and titanium head plates obtained both (K) underneath the head plate material and (L) in the imaging area. (M) CT images with corresponding isodose lines for mice with no head plate, carbon fiber head plate, and titanium head plate.

### TBI treatments, head plate dosimetry, and dose simulations

We used the Precision X-RAD SMART Plus / 225cx small animal image guided irradiation system (Precision X-Ray, North Branford CT USA) (24) to administer TBI treatments in a single-fraction of a 2 or 10 Gy soft-tissue-equivalent dose, acquire CT images, and perform dose calculations (25,26). Further description of treatment and dosimetric evaluation can be found in Appendix 1:E

### Histology Preparation and Scoring

Details of histology preparation can be found in Appendix 1:F

### Statistical analysis

We performed all statistical testing using Prism (V.9.00 (121), GraphPad). For analysis of the treatment effects, a two-way mixed-effects model was performed followed by Tukey’s post-hoc comparison when appropriate. We performed post-hoc comparisons with 3 families and 3 comparisons per family for both time and treatment group comparisons. Measurements of leukemia in the PB of mice in treatment groupswere compared using a one-way mixed-effect model for cross group and longitudinal comparisons. Unless stated otherwise, we performed all other significance testing using a Welch’s two-sided t-test. All distribution error bars are displayed as the mean plus or minus one standard deviation.

## Results

### Study timing and L-QMPM validation

We performed treatment intervention for mice bearing AML and ALLapproximately when the percentage of leukemia in the PB reached the level where changes in BMV function were observed (Figure 1C-D, Table E1-2). An additional group of mice bearing ALL were treated at higher ALL burden to observe the effects of TBI after ALL had caused significant reductions in WIS_tissue_ (Table E3)

To understand the differences between the timing of single-vessel blood flow changes in mice bearing AML or ALL, we characterized the growth kinetics of leukemia in the PB, femur, and calvarium. No significant time-matched differences in the percentage of leukemia in the PB of mice were found between mice bearing ALL and AML post leukemia-cell injection (Figure E1 A). However, in the calvarium, we observed significantly higher engraftment of ALL compared to AML (Figure E1 B-C). Similar differences were found in the femur bone marrow (Figure E1 C-D). The data suggest that differences in vascular function may be due to differences in the onset of leukemia engraftment in the calvarium and femur bone marrow.

We observed no significant differences in any imaging parameters in WT untreated mice between imaging time-points (Table E4). Additionally, no significant differences in any of the imaging parameters or leukemia PB measurements were observed prior to TBI treatments in treatment subgroups with matching types of disease, validating longitudinal and cross-group comparisons post-treatment (Table E1-2). Leukemia growth and treatment response could be observed in the calvarium using L-QMPM (Figure 1E-I Figure E2).

We performed dosimetry measurements at different regions underneath titanium and carbon fiber head plates, as well as performed CT-based dose simulations to ensure accurate dose delivery and dose evaluation of the calvarium after cranial window surgery, respectively. Significant differences in dose were noticed between a lack of head plate and directly under the titanium head plate material, whereas no significant differences were noticed between a lack of head plate and carbon fiber headplates (Figure 1J-L). We observed significant artificats in CT images ofmice with titanium head plates. However, noartifacts were present for carbon fiber head plates. (Figure 1M). For these reasons, we utilized carbon fiber head plates for all L-QMPM imaging studies.

### AML and ALL increased vessel density, reduced blood flow, and reduced WIS_tissue_

We split L-QMPM imaging data into relative low and high leukemic burden groups using PB measurements based on the timing of BMV changes. Reduced vascular diameter, increased vessel density, and reduced single-vessel blood flow were observed in mice bearing leukemia at both low and high leukemic burden compared to WT mice (Figure 2A-D, Table E5).

**Figure 2:**
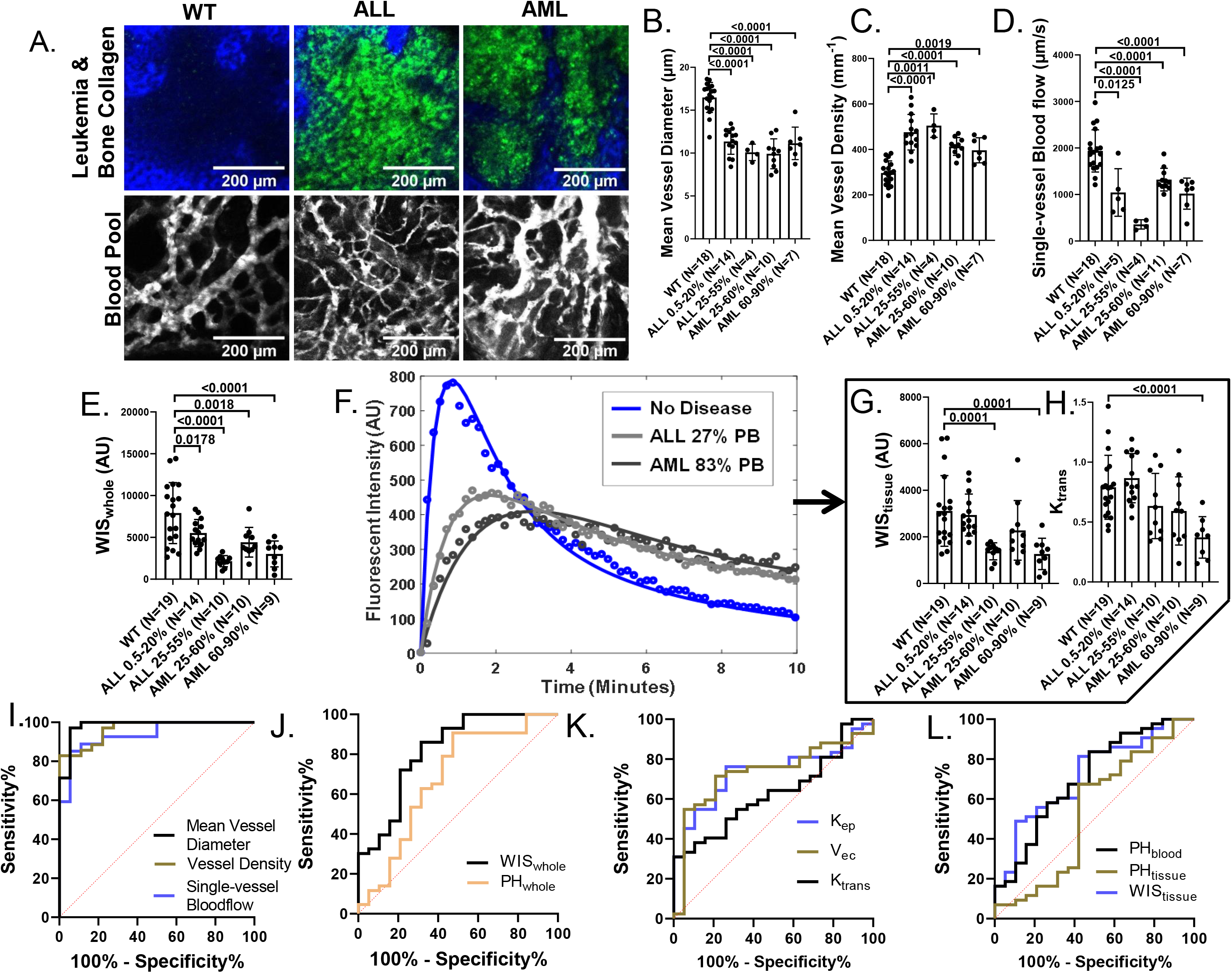
BMV remodeling with the onset of leukemia. (A) Top: images of GFP+ leukemia (green) and second harmonic generation from the collagen in the bone (blue); bottom: Images of BMV blood pool fluorescence from Qtracker™ 655 vascular labels (white). L-QMPM measurements of (B) mean vessel diameter, (C) mean vessel density, (D) single-vessel blood flow, (E) WIS_whole_, (G) WIS_tissue_, and (H) K_trans_ for WT mice and mice with varying leukemic burdens measured through PB sampling. (F) Dextran fluorescence from extravascular tissue ROIs. Solid lines indicate the corresponding fitting function used for compartmental modeling. (I-L) Receiver operating characteristic curve for a variety of L-QMPM parameters. A single measurement time-point from each mouse, typically matching the highest untreated disease burden was used for analysis of all imaging parameters.

BMV changes at low disease burden coincided with decreased WIS_whole_ for mice bearing leukemia compared to WT mice (P=0.0178 and P=0.0018, respectively, Figure 2E, Table E5). we observed decreases in K_ep_ and increases in ν_ec_ in mice bearing AML at low leukemic burden compared to WT mice (P<0.0001 and P=0.0412, respectively Table E5).

At high leukemic burden, we observed decreases in WIS_tissue_ for mice bearing ALL and AML compared to WT mice (P=0.0001 and 0.0001, respectively Figure 2F-G, Table E5). Significant differences for time-lapsed imaging parameters in mice bearing leukemia at high disease burden included decreased WIS_tissue_, WIS_whole_, PH_whole_, PH_blood_, K_ep_, as well as increased ν_ec_ compared to WT mice. (Figure 2E-G, Table E5). We additionally observed decreased K_trans_ at high AML burden (P<0.0001, Figure 2H). The most effective time-lapsed imaging parameter to distinguish between mice bearing leukemia and WT mice was WIS_whole_ (Figure 2I-L, Figure E3).

We measured the percent signal contribution to PH_whole_ from the extravascular tissue ROI, to better understand peak height measurments in DCE imaging. Mean signal contribution was 55.7±16.7%, 64.9±3.7%, and 71.7±20.7% for untreated WT mice, and mice bearing ALL or AML, respectively (Figure E4 A). We observed no correlation between PH_whole_ and vascular density using data from untreated WT mice and mice bearing AML and ALL. (R^2^=0.005, P=0.5235, Figure E4B).

### TBI reduced vascular density, dilated blood vessels, and, in mice bearing leukemia, increased blood perfusion

We observed increases in mean vessel diameter, for all mice 2 days after 2 Gy and 10 Gy TBI as well as 5 days after 10 Gy TBI compared to that in pretreatment time-points and untreated mice (Figure 3A-B, Table E1-E4). For mice bearing AML or ALL, we observed increases in single-vessel blood flow 2 days after 2 Gy TBI and 10 Gy TBI compared to levels in pretreatment time-points and untreated mice (Figure 3C, Table E1-E3). Compared to untreated mice, increases in single-vessel blood flow were observed 5 days after 10 Gy TBI treatments for mice bearing leukemia (Figure 3C, Table E1-E2).

**Figure 3:**
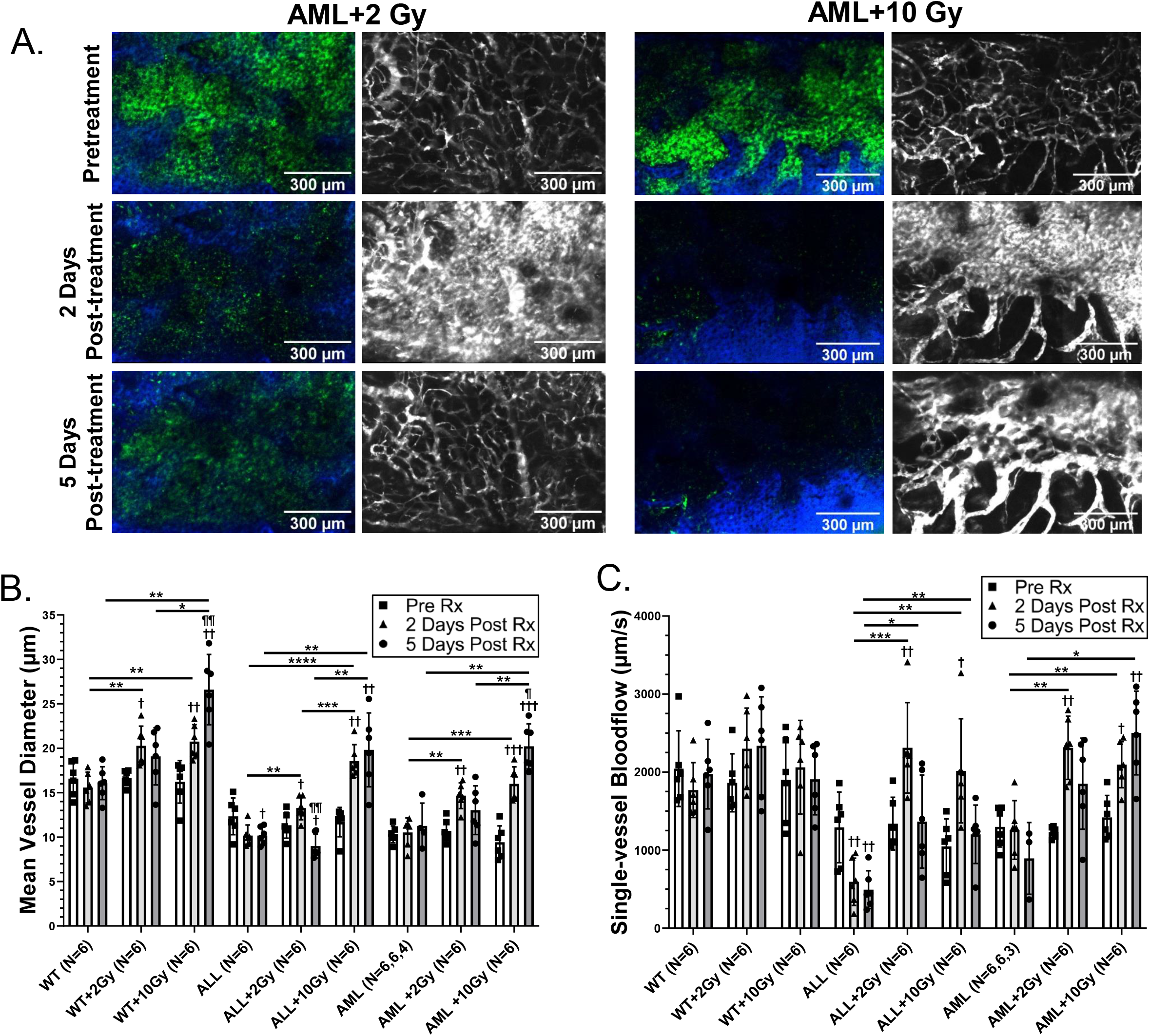
TBI induced changes in leukemic burden, vascular leakiness, single-vessel blood flow, and vessel diameter. (A) Left: images of AML (Green) and second harmonic generation from the collagen in the bone (blue); right: maximum intensity projections of Qtracker™ 655 vascular label (white). Images of 2 Gy and 10 Gy TBI are shown depicting substantial vascular leakage and reduction in AML burden 2 days after TBI. The effects of TBI on (B) mean vessel diameter and (C) single-vessel blood flow for WT mice, and mice bearing ALL and AML (*P < .05, **P < .01, ***P < .001, ****P < .0001, compared to pretreatment time-point: †P < .05, ††P < .01, †††P < .001, ††††P < .0001, compared to 2 days post-treatment time-point ¶P < .05, ¶¶P < .01, ¶¶¶P < .001, ¶¶¶¶P < .0001).

We observed decreases in vessel density 2 days after 10 Gy TBI for mice bearing leukemia, compared to that in pretreatment time-points and, for mice bearing ALL, untreated mice (Figure E4C). Decreases in vessel density were also observed 5 days after 10 Gy TBI for all mice, compared to levels in pretreatment time-points and untreated mice. Decreases in vessel density and increases in vessel diameter were also observed 5 days after 10 Gy TBI compared to 2 Gy for all mice. We observed similar trends between femur histology, calvarium histology, and L-QMPM imaging of the BMV for vessel diameter and vessel density measurements (Figure 4A-D).

**Figure 4:**
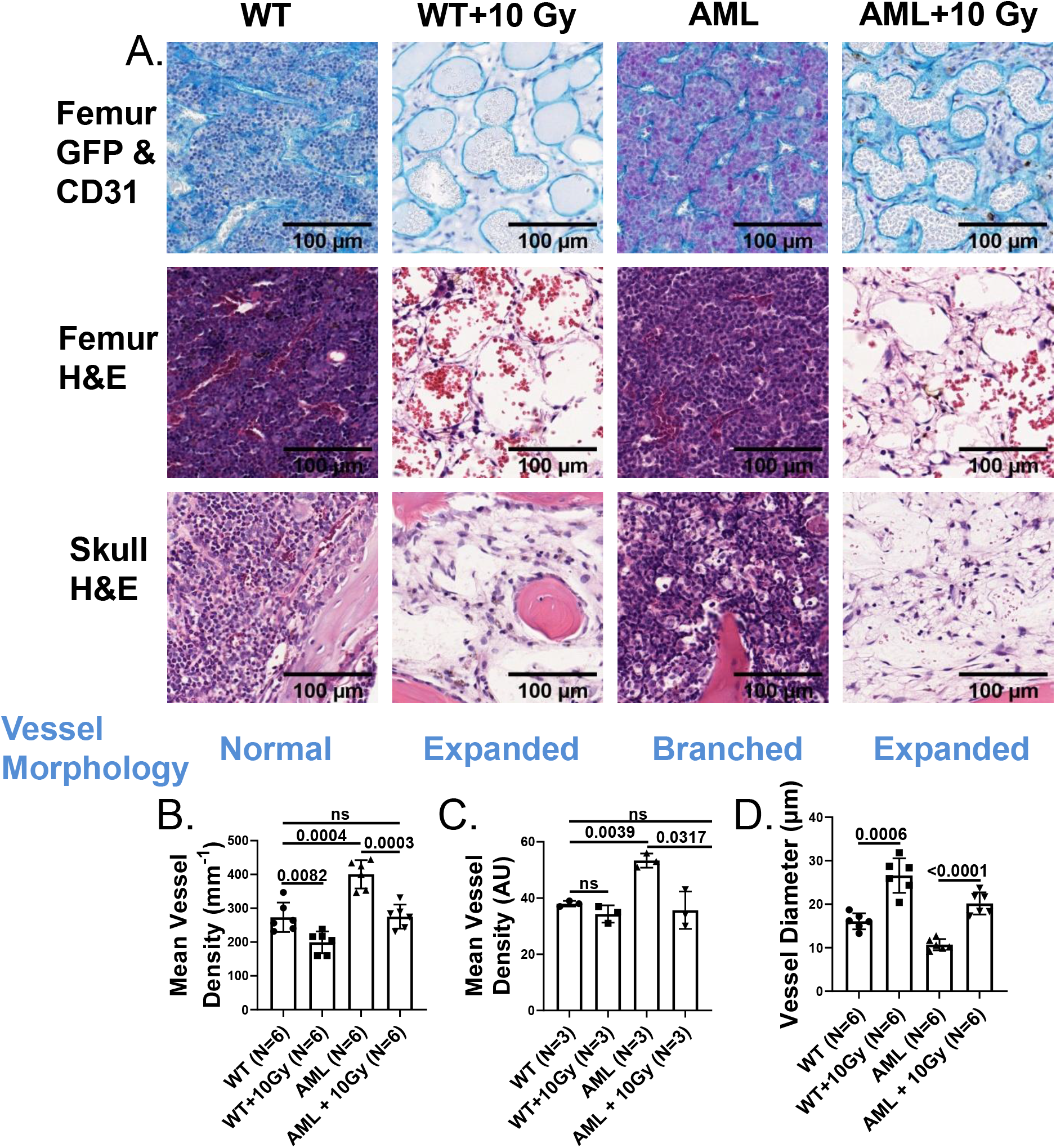
Histological validation of L-QMPM imaging. **(A)** Top: Femur histology sections stained for GFP+ AML cells (purple) and CD31 endothelial cells (blue) (top); middle: femur hematoxylin and eosin stained sections (middle); bottom: skull stained for hematoxylin and eosin sections (bottom). Vessel dilation was observed 5 days after 10Gy TBI for WT mice and mice bearing AML. (B) Mean vessel density measurements from L-QMPM imaging of mice with matching conditions to histology. (C) Vessel density scoring of individual vessels in CD31+ endothelial stained femur sections. Two different image locations were used for vessel scoring. (D) Vessel diameter measurements from L-QMPM imaging of mice with matching treatment conditions to histology.

### Radiotherapy increased drug delivery potential to the healthy and leukemic bone marrow

We observed increases in WIS_tissue_ for all mice 2 days after both 2 Gy TBI and 10 Gy TBI compared to either pretreatment time-points, or untreated mice (Figure 5 A-D, Table E1-4). Percent increases in WIS_tissue_ mean values ranged from 39% to 81% for WT mice and 139% to 227% for mice bearing leukemia compared to pretreatment. Similar increases post-treatment for K_trans_ and K_ep_ were observed (Figure 5E, Table E1-4). We observed increases in WIS_tissue_ 5 days after 10 Gy TBI for mice bearing leukemia compared to untreated mice, and, for mice bearing ALL, pretreatment time-points. We additionally observed significant increases in WIS_tissue_ 5 days after 10 Gy TBI compared to 2Gy for mice bearing AML or ALL (mean percent increases of 165% and 188%, respectively). We found significant decreases in WIS_tissue_ prior to treatment intervention in mice bearing AML or mice bearing ALL treated with 10 Gy at high disease burden compared to WT mice (P=0.0279 and 0.0003, respectively, Figure E4D), demonstrating that increases in WIS_tissue_ after TBI occurred in healthy or leukemic bone marrow microenvironments. We observed increases in PH_tissue_ and ν_ec_ after TBI at a variety of doses and time-points likely due to changes in cellularity (Table E1-4). Similar results are observed in DCE imaging (27).

**Figure 5:**
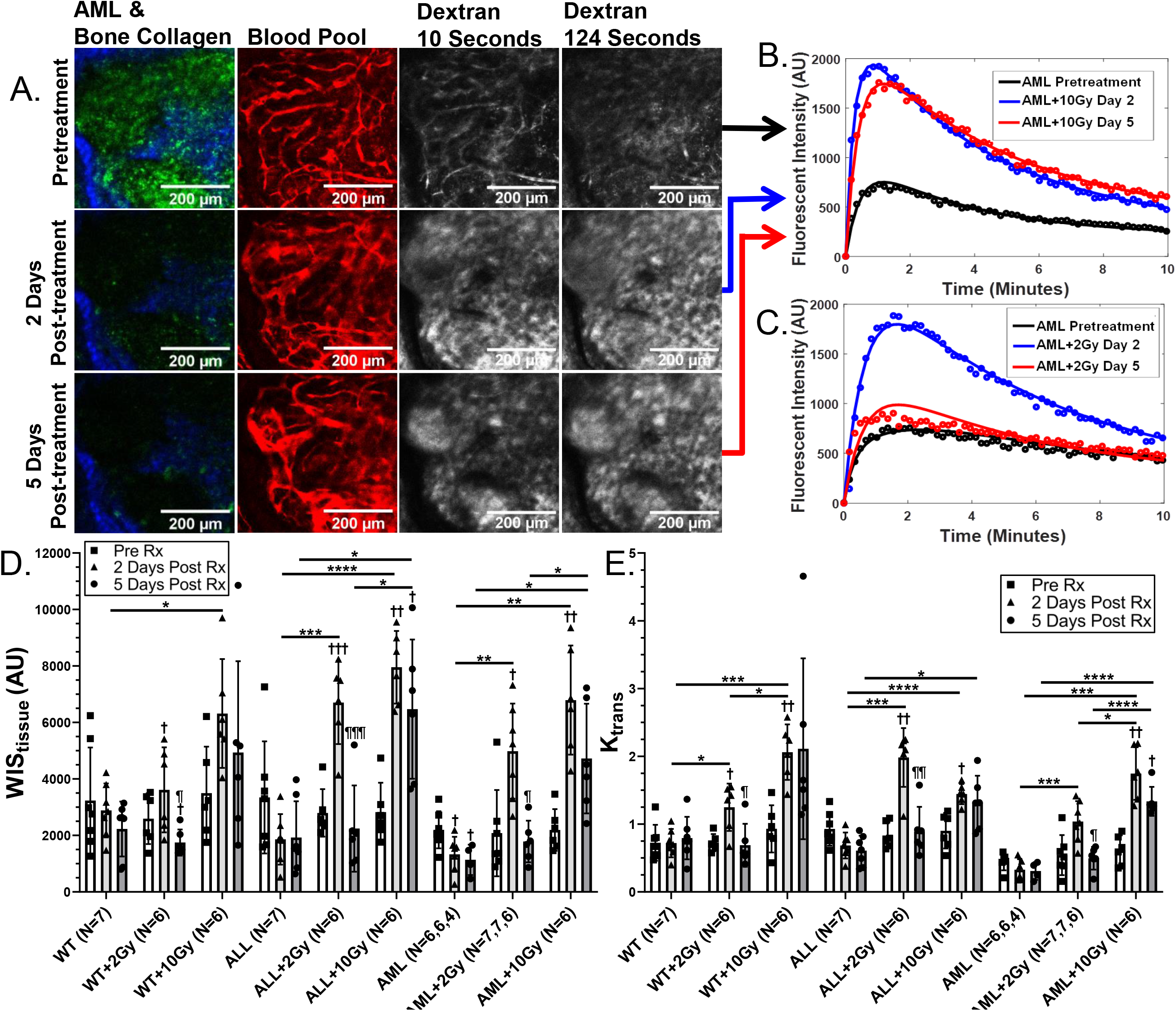
Measurements of BMV drug delivery potential. (A) Images of AML (green) and second harmonic generation from the collagen in the bone (blue) (far left). Images of Qtracker™ 655 vascular label (red) (left middle). Images of of dextran (white) from the first frame after injection of dextran (right middle). Images of dextran (white) from the 12^th^ frame after dextran injection (far right). A dose of 10 Gy TBI was given to the imaged mouse. (B) Fluorescent intensity values from the extravascular tissue ROI taken from time-lapsed images of dextran from the mouse images shown in Figure 5A. Solid lines indicate the extravascular fitting function for compartment modeling. (C) Fluorescent intensity values from the extravascular tissue ROI taken from a representative mouse treated with 2 Gy TBI. The effects of TBI on (D) WIS_tissue_, and (E) K_trans_ for WT mice and mice bearing ALL or AML. (*P < .05, **P < .01, ***P < .001, ****P < .0001, compared to pretreatment time-point: †P < .05, ††P < .01, †††P < .001, ††††P < .0001, compared to 2 days post-treatment time-point ¶P < .05, ¶¶P < .01, ¶¶¶P < .001, ¶¶¶¶P < .0001 Tukey’s post hoc comparison).

A summary of the effects of TBI is shown in Figure 6A. We found positive linear correlations with WIS_tissue_ for K_trans_, K_ep_, ν_ec_, single-vessel blood flow, and mean vessel diameter, while no correlation was found for vessel density (Figure 6B-G).

**Figure 6:**
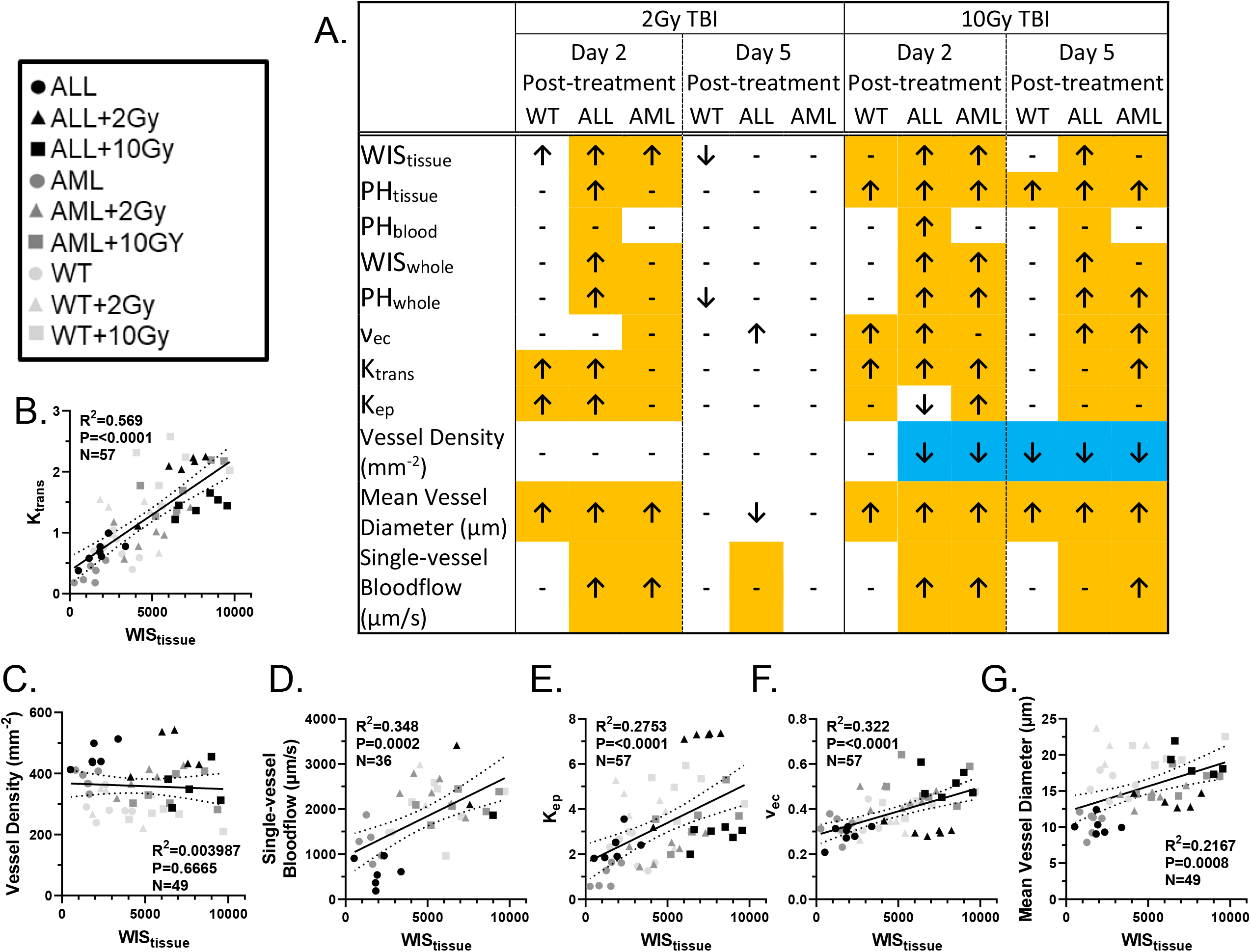
Summary of TBI treatment effects, and correlation to WIS_tissue_. (A) A summary of the response of WT mice, and mice bearing ALL and AML to 2 Gy and 10 Gy TBI treatments. Arrows indicate a significant increase or decrease compared to pretreatment time-points, orange indicates a significant increase compared to the untreated group, and blue indicates a decrease compared to the untreated group (P<0.05 for all). (B-G) Linear correlation plots for various L-QMPM imaging parameters plotted with data from the time-point 2 days after TBI treatments. A single image time-point was used per mouse for linear correlation calculations.

## Discussion

We developed L-QMPM imaging to measure several parameters that quantified the BMV and drug delivery potential before and after radiotherapy. Decreased WIS_tissue_, WIS_whole_, PH_whole_, K_ep_, single-vessel blood flow and mean vessel diameter, as well as increased vessel density and ν_ec_ were observed with the onset of leukemia. WIS_whole_ was the most effective contrast-based, time-lapsed imaging parameter to identify mice bearing leukemia. As hypothesized, 2 Gy and 10 Gy TBI treatments increased WIS_tissue_, K_trans_, and mean vessel diameter in WT mice and mice bearing leukemia 2 days after TBI. We also observerd increased single-vessel blood flow for mice bearing AML and ALL. Changes in BMV morphology and function were observed 5 days after 10 Gy TBI, while 2 Gy caused minimal changes. Linear correlations between WIS_tissue_ and vessel diameter, single-vessel blood flow, K_trans_, ν_ec_ and K_ep_ were found, while vessel density was not correlated with WIS_tissue_.

DCE imaging analysis applies mathematical models to or infers the BMV changes that influence the delivery of contrast agents. Wee calculated WIS_whole_ and PH_whole_ as measurements commonly interpreted as the volumetric tissue blood flow rate, and tissue blood volume for DCE imaging (8). Several studies using DCE imaging identified changes in PH_whole_ or similar mathematically modeled parameters to be positively correlated with vessel density and have proposed DCE imaging as an alternative method to quantify vessel density in the untreated malignant bone marrow (10,28). This circumstance is desirable as it would reduce the need for invasive bone marrow biopsies to quantify vessel density, which is a prognostic marker for tumor aggressiveness (29). However in this work, we found no correlation between PH_whole_ and vessel density with leukemia onset likely due to the reduction in mean vessel diameter present in mice bearing leukemia. Additionally, over half of the fluorescent signal contributing to PH_whole_ inwas from the extravascular ROI, suggesting that PH_whole_ measurements may be significantly influenced by leakage of contrast into the extravascular tissue. Our results demonstrate that positive correlations of PH_whole_ with vessel density or vessel volume cannot be assumed without pathological validation, and that mathematical modeling should be used to account for signal contributions from the extravascular tissue when quantifying vascular parameters using DCE.

We identified WIS_whole_ as the most effective time-lapsed imaging measurement to identify mice bearing leukemia. As WIS_whole_ is primarily a measurement of volumetric blood flow, it will be closely related to mean vessel diameter, vessel density, and single-vessel blood flow, which are all affected early in leukemia progression. This enables WIS_whole_ to better identify mice bearing leukemia than other contrast-based time-lapsed imaging parameters, which are less sensitive to early vascular alterations.

Because WIS_whole_ is primarily a measurement of volumetric blood flow, it is enticing to infer improved drug delivery potential when increases in WIS_whole_ are observed. However, we observed that changes in WIS_whole_ and WIS_tissue_ did not occur at the same disease burden. Additionally, changes in WIS_tissue_ were observed in WT mice after TBI, but no changes were observed in WIS_whole_. Our results demonstrated that WIS_whole_ may not accurately identify changes in drug delivery potential, likely because WIS_whole_ is a convolution of signals from vascular and extravascular tissues. These observations identified the importance of validating drug delivery potential when changes in DCE imaging parameters are observed.

Increases in WIS_tissue_ 2 days after TBI in WT mice were accompanied only by increases in K_trans_ and mean vessel diameter, suggesting that increases in WIS_tissue_ were due to increased vascular permeability or increases in vascular surface area, which commonly influence K_trans_ (23). Similar alterations in K_trans_ after TBI have been observed previously using DCE-MRI (27). For mice bearing leukemia, increases in blood flow after TBI may also contribute to increased WIS_tissue_. Similar observations of improved tissue drug perfusion after radiotherapy have been made in solid tumor models (30,31), suggesting that the leukemic BMV and solid tumor vasculature may have similar responses to radiotherapy. Our results suggest that increased WIS_tissue_ may be found shortly after a variety of TBI doses, and directly demonstrate the role of the BMV in previous observations of improved cellular chemotherapy uptake after neo-adjuvant radiotherapy (12).

Although we observed similar changes in WIS_tissue_ 2 days after 2 Gy and 10 Gy TBI, the duration of their changes were different. This finding suggests that the window of opportunity for synergistic combination therapy with therapeutics after reduced-intensity TBI regimens may be smaller than with standard myeloblative regimens of TBI (32). We observed significant changes in vessel density and vessel diameter 5 days after 10 Gy TBI compared to 2 days, while no significant differences were noticed in WIS_tissue_. These data suggest that maximum BMV morphological changes after TBI may occur later than the maximum increases in WIS_tissue_. These findings are in agreement with other studies observing vasodilation and macromolecular contrast leakage 4-7 days after myeloblative doses (33,34). Future studies should identify how long-term vascular damage after myeloblative TBI and bone marrow transplantation influence drug delivery potential.

We observed decreases in vessel density 2 days after 10 Gy treatment for mice bearing leukemia, but not for WT mice, suggesting that the leukemic BMV may have increased radiosensitivity. Newly formed immature vessels have been shown to be sensitive to radiotherapy (35,36) and more mature vascular structures have been found after fractionated radiotherapy in solid tumors (37). This suggests that the appropriate radiotherapy treatment may be able to preserve mature vessels, while eliminating newly formed vessels present from leukemia signaling. Synergistic effects have been observed with radiotherapy and vascular based therapeutic agents for solid tumors (38), making combination therapy an attractive option for the treatment of leukemia. Future studies employing dose fractionation need to be performed to understand how radiotherapy can be used to normalize the vascular system, and whether enhancements to drug delivery potential are observed after successive fractions of low dose radiotherapy

Although L-QMPM is useful to accurately assess drug delivery potential, it is limited to preclinical models, making DCE imaging modalities such as MRI or CT necessary for clinical translation. In this work, we provide direct observations of BMV morphology and function, as well as measurements used in DCE imaging. These observations will be invaluable for interpretation of DCE imaging of the bone marrow.

## Conclusion

We identified WIS_whole_ as the most effective parameter to distinguish between WT mice and mice bearing AML or ALL. This suggests that DCE imaging should focus on the rate of initial increase in whole tissue contrast signal during time-lapsed imaging as a biomarker for leukemia. Our results show that the response of the leukemic BMV to neo-adjuvant radiotherapy improves drug delivery to the bone marrow by improving blood flow and increasing drug permeation through the BMV to the extravascular tissue.

## References

1. Passaro D, Di Tullio A, Abarrategi A, et al. Increased vascular permeability in the bone marrow microenvironment contributes to disease progression and drug response in acute myeloid leukemia. Cancer Cell 2017;32:324–341.e6.

2. Duarte D, Hawkins ED, Akinduro O, et al. Inhibition of endosteal vascular niche remodeling rescues hematopoietic stem cell loss in aml. Cell Stem Cell.

3. Hussong JW, Rodgers GM, Shami PJ. Evidence of increased angiogenesis in patients with acute myeloid leukemia. Blood 2000;95:309–313.

4. Schaefer C, Krause M, Fuhrhop I, et al. Time-course-dependent microvascular alterations in a model of myeloid leukemia in vivo. Leukemia 2007;22:59.

5. Ossenkoppele GJ, Stussi G, Maertens J, et al. Addition of bevacizumab to chemotherapy in acute myeloid leukemia at older age: A randomized phase 2 trial of the dutch-belgian cooperative trial group for hemato-oncology (hovon) and the swiss group for clinical cancer research (sakk). Blood 2012;120:4706–4711.

6. Jain P, Lee HJ, Qiao W, et al. Fcr and bevacizumab treatment in patients with relapsed chronic lymphocytic leukemia. Cancer 2014;120:3494–501.

7. Haibe Y, Kreidieh M, El Hajj H, et al. Resistance mechanisms to anti-angiogenic therapies in cancer. Frontiers in Oncology 2020;10.

8. Cuenod CA, Balvay D. Perfusion and vascular permeability: Basic concepts and measurement in dce-ct and dce-mri. Diagnostic and Interventional Imaging 2013;94:1187–1204.

9. Faye N, Fournier L, Balvay D, et al. Dynamic contrast enhanced optical imaging of capillary leakage. Technology in Cancer Research & Treatment 2011;10:49–57.

10. Shih TTF, Tien HF, Liu CY, et al. Functional mr imaging of tumor angiogenesis predicts outcome of patients with acute myeloid leukemia. Leukemia 2006;20:357–362.

11. Petrakis NL, Masouredis SP, Miller P, et al. The local blood flow in human bone marrow in leukemia and neoplastic diseases as determined by the clearance rate of radioiodide (i131). The Journal of Clinical Investigation 1953;32:952–963.

12. Brooks J, Kumar B, Zuro DM, et al. Biophysical characterization of the leukemic bone marrow vasculature reveals benefits of neoadjuvant low-dose radiation therapy. International Journal of Radiation Oncology, Biology, Physics 2021;109:60–72.

13. Jung Y, Spencer JA, Raphael AP, et al. Intravital imaging of mouse bone marrow: Hemodynamics and vascular permeability. In: Ishii M, editor Intravital imaging of dynamic bone and immune systems: Methods and protocols. New York, NY: Springer New York; 2018. pp. 11–22.

14. Dombret H, Gardin C. An update of current treatments for adult acute myeloid leukemia. Blood 2016;127:53–61.

15. Pustylnikov S, Sagar D, Jain P, et al. Targeting the c-type lectins-mediated host-pathogen interactions with dextran. J Pharm Pharm Sci 2014;17:371–392.

16. Dewhirst MW, Secomb TW. Transport of drugs from blood vessels to tumour tissue. Nat Rev Cancer 2017;17:738–750.

17. Manlove LS, Berquam-Vrieze KE, Pauken KE, et al. Adaptive immunity to leukemia is inhibited by cross-reactive induced regulatory t cells. Journal of immunology (Baltimore, Md: 1950) 2015;195:4028–4037.

18. Hossain DMS, Dos Santos C, Zhang Q, et al. Leukemia cell–targeted stat3 silencing and tlr9 triggering generate systemic antitumor immunity. Blood 2014;123:15–25.

19. Egawa G, Ono S, Kabashima K. Intravital imaging of vascular permeability by two-photon microscopy. In: Nagamoto-Combs K, editor Animal models of allergic disease: Methods and protocols. New York, NY: Springer US; 2021. pp. 151–157.

20. Schindelin J, Arganda-Carreras I, Frise E, et al. Fiji: An open-source platform for biological-image analysis. Nature Methods 2012;9:676.

21. Preibisch S, Saalfeld S, Tomancak P. Globally optimal stitching of tiled 3d microscopic image acquisitions. Bioinformatics 2009;25:1463–1465.

22. Otsu N. A threshold selection method from gray-level histograms. IEEE Transactions on Systems, Man, and Cybernetics 1979;9:62–66.

23. Tofts PS. Modeling tracer kinetics in dynamic gd-dtpa mr imaging. Journal of Magnetic Resonance Imaging 1997;7:91–101.

24. van Hoof SJ, Granton PV, Verhaegen F. Development and validation of a treatment planning system for small animal radiotherapy: Smart-plan. Radiotherapy and oncology: journal of the European Society for Therapeutic Radiology and Oncology 2013;109:361–6.

25. Faddegon BA, Kawrakow I, Kubyshin Y, et al. The accuracy of egsnrc, geant4 and penelope monte carlo systems for the simulation of electron scatter in external beam radiotherapy. Physics in medicine and biology 2009;54:6151–63.

26. Downes P, Jarvis R, Radu E, et al. Monte carlo simulation and patient dosimetry for a kilovoltage cone-beam ct unit. Medical physics 2009;36:4156–67.

27. Wang K, Zha Y, Lei H, et al. Mri study on the changes of bone marrow microvascular permeability and fat content after total-body x-ray irradiation. Radiation research 2018;189:205–212.

28. Moehler TM, Hawighorst H, Neben K, et al. Bone marrow microcirculation analysis in multiple myeloma by contrast-enhanced dynamic magnetic resonance imaging. International Journal of Cancer 2001;93:862–868.

29. Hlatky L, Hahnfeldt P, Folkman J. Clinical application of antiangiogenic therapy: Microvessel density, what it does and doesn’t tell us. JNCI: Journal of the National Cancer Institute 2002;94:883–893.

30. Potiron VA, Abderrahmani R, Clément-Colmou K, et al. Improved functionality of the vasculature during conventionally fractionated radiation therapy of prostate cancer. PloS one 2013;8:e84076–e84076.

31. Kleibeuker EA, Fokas E, Allen PD, et al. Low dose angiostatic treatment counteracts radiotherapy-induced tumor perfusion and enhances the anti-tumor effect. Oncotarget 2016;7:76613–76627.

32. Gyurkocza B, Sandmaier BM. Conditioning regimens for hematopoietic cell transplantation: One size does not fit all. Blood 2014;124:344–353.

33. Le V-H, Lee S, Lee S, et al. In vivo longitudinal visualization of bone marrow engraftment process in mouse calvaria using two-photon microscopy. Scientific Reports 2017;7:44097.

34. Slayton WB, Li X-M, Butler J, et al. The role of the donor in the repair of the marrow vascular niche following hematopoietic stem cell transplant. STEM CELLS 2007;25:2945–2955.

35. Grabham P, Hu B, Sharma P, et al. Effects of ionizing radiation on three-dimensional human vessel models: Differential effects according to radiation quality and cellular development. Radiation research 2011;175:21–8.

36. Park M-T, Oh E-T, Song M-J, et al. The radiosensitivity of endothelial cells isolated from human breast cancer and normal tissue in vitro. Microvascular Research 2012;84:140–148.

37. Chen F-H, Fu S-Y, Yang Y-C, et al. Combination of vessel-targeting agents and fractionated radiation therapy: The role of the sdf-1/cxcr4 pathway. International Journal of Radiation Oncology, Biology, Physics 2013;86:777–784.

38. Wachsberger P, Burd R, Dicker AP. Tumor response to ionizing radiation combined with antiangiogenesis or vascular targeting agents. Exploring Mechanisms of Interaction 2003;9:1957–1971.

