## Supplemental Materials for "Longitudinal preclinical imaging characterization of drug delivery potential after radiotherapy in the healthy and leukemic bone marrow vascular microenvironment"

**Supplementary text**

#### *Appendix A: Mice and cell injections & study timing*

For initial non-imaging studies of ALL growth kinetics, 6-week-old C57BL/6N mice (Strain 556, Charles River, Wilmington, MA) were used. Initial growth kinetic studies of AML were performed with 9-week-old male and female albino B6 mice (B6 Tyrc-2J/J-B6 stockNo: 000058, Jackson Laboratories, Bar Harbor, ME). For ALL and AML cell injections, cells were suspended in 200 µl or 100 µl of phosphate buffered saline (PBS) respectively. ALL and AML injections were administered using tail vein or retro-orbital injections respectively. Each treatment group for healthy and leukemic mice consisted of a minimum of two mice of each sex.

Surgery on WT mice was performed 2 days before the start of imaging. ALL injections and headplate surgery on mice bearing ALL were performed 7-8 days and 2-3 before the start of imaging respectively. ALL was detectable in the peripheral blood (PB) of all sampled mice at the first imaging timepoint with PB readings ranging from 0.5% to 12.9%. Approximately 75% of the mice bearing ALL were sampled. An additional group of mice bearing ALL were treated with 10Gy at higher burden. ALL injections and headplate surgery for these mice was performed 10 and 2 days before the start of imaging respectively. PB readings of these mice at the start of imaging ranged from 32% to 55% for all mice. Headplate surgery on mice bearing AML was performed after mice reached 25% AML cells in PB. AML injections and headplate surgery were performed 10-16 and 2 days before the start of imaging, respectively. AML in the PB at the start of imaging ranged from 28% to 75.8% for all mice bearing AML.

#### *Appendix B: Cranial window surgery*

Two days before surgery mice were given MediGel® Sucralose gel cups (Clear $H_2O$, Portland, ME) mixed with 1.125 mg of carprofen per cup. An additional cup was given on the day of surgery. For surgery, mice were anesthetized using 5% isoflurane initially at a flow rate of 1.5 L/min, and then maintained with 1.3% isoflurane at a flow rate of 0.8 liters per minute. A stereotactic apparatus with a bite bar was utilized to keep the mouse stable during surgery. The hair on the top of the skull was removed using a hair trimmer and the skin above the skull was sterilized using a topical antiseptic (Betadine®, Purdue Products L.P., Stamford, CT), and rinsed with saline. An incision was made to remove the skin and membrane above the calvarium, and a custom head plates made of carbon fiber or titanium with an inner diameter of 7mm or 8mm respectively was fixed to the calvarium using glass ionomer cement (Pearson PQ, Sylmar, CA) (Figure 1A). A round 5mm glass coverslip was placed on top of the exposed skull in the middle of the headplate, and was sealed with cyanoacrylate glue (Loctite 401™, Henkel, Düsseldorf, Germany). Mice were administered 0.5-1.0 mg/kg of buprenorphene-SR (ZooPharm, Fort Collins, CO) as an additional analgesic at the time of surgery.

#### *Appendix C: QMPM image acquisition and analysis*

Fluorescent excitation for the Prairie Ultima multiphoton microscope was performed using a Chameleon Ultra II tunable Ti:Sapphire laser with 140 femtosecond pulses (Coherent, Santa Clara, CA). An Olympus XLUMPlanFL 20x objective (1.00 NA water objective) was used for image acquisition of all images. Simultaneous four channel acquisition was performed using 660/40 (far red), 595/50 (red), 525/50 (green), and 460/50 (blue) filters for visualization of GFP+ leukemia, blood pool fluorescent agents and bone-collagen. Data acquisition using the Prairie Ultima microscope was handled by Prairieview 5.3 software and imaging was performed at room temperature.

For imaging, mice were anesthetized using 5% isoflurane initially at a flow rate of 1.5 L/min, and then maintained with 1.3% isoflurane at a flow rate of 0.8 liter per minute. The stereotactic head plate was inserted into a custom-built heated stage maintained at 37°C allowing for easy viewing of the mouse calvarium. Imaging was performed on the frontal bone region of the calvarium, near the sagittal suture for all mice. Mice were catheterized intravenously through the tail vein and injected with 350 µg of 4kDa TRITC dextran suspended in 100 µl of phosphate buffered saline (PBS) during time-lapsed imaging to observe drug delivery potential. TRITC dextran fluorescent excitation was performed at 820 nm. Time-lapsed imaging consisted of 120 frames with 10.3 second frame interval at a 512 by 512 pixel resolution. To identify the vascular region from the tissue region, a second injection of either 400 µg of 2mDa FITC dextran suspended in 100 µl of PBS or 10µl of Qtracker™ 655 Vascular Label mixed with 90 µl of PBS was performed. Immediately after injection, a series of fast, five-slice z-stack images with 15 µm spacing, 512 by 512 pixel resolution, and no averaging were taken before tissue leakage of FITC dextran or Qtracker™ occurred. Substantial leakage was often present in mice treated with TBI due to loss in cellularity and vascular damage. This made clear viewing vascular morphology difficult after the first 2-3 minutes in some treated mice. Three separate fast z-stack images were used to manually measure mean vessel density and mean vessel diameter for each mouse using Fiji/Imagej. FITC dextran excitation was performed at 820nm. Qtracker™ excitation, GFP+ leukemia excitation, and second harmonic generation of the collagen in the bone was performed at 900nm. Tiled z-stack images of ALL and blood pool fluorescent contrast with 10um slice spacing were obtained by overlapping individual images by 15-20% with a 512 by 512 pixel resolution and 2x averaging.

#### *Appendix D: Image segmentation and time-lapsed imaging analysis*

To identify the vascular tissue ROI, an Otsu's threshold was applied to blood pool agent images of the vasculature. To identify the extravascular tissue ROI, the vascular tissue ROI was cropped out of the 4kDa dextran image and an Otsu's threshold was applied to the 15$^{th}$ frame of the dextran time-lapsed images. If a single threshold was not adequate to properly segment the extravascular tissue ROI from the background, the first extravascular ROI region was subtracted from the image and a second Otsu's threshold was applied. The two extravascular tissue regions were then combined to identify the extravascular tissue ROI. Two thresholds were only applied in a few cases where there was extensive heterogeneity in tissue signal intensity in the image. The whole tissue ROI was identified by combining extravascular and vascular ROIs. After identifying initial ROIs, boundary edge erosion was applied to the regions to better isolate dextran fluorescent signal coming from the respective tissue compartments. An 8 pixel boundary erosion was applied to extravascular and whole tissue ROIs. A 4 pixel boundary erosion was applied to the vascular tissue ROI.

Time-lapsed imaging of 4kDa TRITC dextran were analyzed using a custom Matlab® (R2018a 9.41.0.81364, MathWorks Natick, MA) script obtaining compartmental modeling parameters and descriptive curve analysis of dextran uptake. To do this, images frames were co-registered to minimize spatial drift between frames of time-lapsed images. For calculation of $K_{trans}$, $K_{ep}$, and $\nu_{ec}$ removal of saturated pixels from extravascular and vascular ROI's was performed prior to analysis. To do this, the percentage of saturated pixels from the ROI with the highest percentage of saturated pixels was identified, and that percentage of pixels was removed from both the vascular and extravascular ROIs. Saturated pixels were only present for a subset of the total images, and in general the removal of saturated pixels made little change to the overall signal intensity from ROIs. The concentration of dextran in the vascular tissue ROI was well described by and fit to a function of two separately weighted bi-exponentials. The concentration of dextran in the extravascular tissue ROI was modeled using the equation.

$$\frac{dC_t(t)}{dt} = (K_{trans})(C_p(t) - \frac{C_t(t)}{v_{ec}})$$

Where $C_t(t)$ and $C_p(t)$ are the concentrations of fluorescent dextran at time t in the extravascular and vascular tissue ROIs respectively. Analysis of the extravascular tissue ROI signal contributions to $PH_{whole}$ was performed using the equation:

$$SC_{tissue} = \frac{\mathrm{SI}_{tissue} * A_{tissue}}{\mathrm{SI}_{tissue} * A_{tissue} + SI_{blood} * A_{blood}}$$

Where $SI_{tissue}$ and $SI_{blood}$ are the mean fluorescent intensities from the extravascular and vascular tissue ROIs after boundary erosion is performed at the time that $PH_{whole}$ is measured. $A_{tissue}$ and $A_{blood}$ are the areas covered by the extravascular and vascular tissue ROIs before boundary erosion is performed.

#### *Appendix E: TBI treatments, CT imaging, and headplate film dosimetry*

TBI treatments for L-QMPM imaged mice were performed while mice were conscious. Treatments were performed with x-ray tube settings of 225 kVp and 13 mA with a 0.32 mm copper filter. CT imaging was performed on mice under anesthesia (2% isoflurane at a flow rate of 1.5 liters per minute) with x-ray tube settings of 40 kVp and 13 mA at a 0.2 mm voxel size with a 2.0 mm aluminum filter. Film dosimetry was performed using Gafchromic™ EBT3 film (Ashland Specialty Ingredients, Bridgewater NJ). The film was initially calibrated at the isocenter of the system after verification of the dose with an externally calibrated ion chamber (PTW TN30013 Farmer Chamber, Freiburg, Germany). An output dose of approximately 1.5 Gy was chosen for headplate dosimetry as it was the dose in the center of the linear response of the film (data not shown).

#### *Appendix F: Histology*

After completion of imaging 5 days after radiotherapy, bone tissues were dissected and fixed in 10% neutral buffered formalin for 48 hours. Bone tissues were decalcified using Richard-Allan Scientific™ Decalcifying Solution (Thermo Scientific™) with 6 hours incubation and raised. Dehydration, clear and paraffinization was performed on a Tissue -Tek VIP Vacuum Infiltration Processor (Sakura Finatek, Torrence, CA, USA). The samples were then embedded in paraffin using a Tissue-Tek TEC Tissue Embedding Station (Sakura Finatek), cut at 5 µm and stained with a Tissue -Tek Prism Plus automated H&E Stainer (Sakura Finatek) according to standard laboratory procedures.

Dual IHC stain for CD31 (Clone: D8V9E Rabbit monoclonal antibody, Cell Signaling Technologies, Danvers, MA, USA) and GFP (Clone: D5.1 Rabbit monoclonal antibody, Cell Signaling Technologies, Danvers, MA) was performed on Ventana Discovery Ultra automated IHC stainer (Ventana Medical Systems, Roche Diagnostics, Indianapolis, USA). Briefly, after deparaffinization, rehydration, endogenous peroxydase activity inhibition and antigen retrieval, the two antigens were sequentially detected, and heat inactivation was performed to prevent antibody cross-reactivity between the same species. Following each primary antibody incubation, DISCOVERY anti-Rabbit HQ and DISCOVERY anti-HQ-HRP were incubated. The stains were then visualized with DISCOVERY Teal kit and DISCOVERY Purple Kit, respectively, counterstained with haematoxylin (Ventana), and coverslipped. Whole slide images were acquired with a Ventana iScan HT Scanner (Roche Diagnostics, Indianapolis, IN, USA) and viewed by iScan image viewer software. Histological vessel density scoring was performed manually using femur sections stained for CD31 to identify endothelial cells. Two separate images were quantified for vessel density per mouse. All pathological findings were verified by a board-certified pathologist with expertise in bone marrow pathology (xx).

#### Table E1

| AML | | Pretreatment | Day 2 Post-treatment | Day 5 Post-treatment |
|---|---|---|---|---|
| | | Mean ± Standard Deviation | Mean ± Standard Deviation | Mean ± Standard Deviation |
| % AML in PB | No RT (N=8,8,4) | 47.7 ± 7.7 | 63.0 ± 12.0† | 82.4 ± 6.7† |
| | 2Gy (N=7) | 54.9 ± 15.5 | # | # |
| | 10Gy (N=6) | 47.9 ± 8.8 | # | # |
| $WIS_{tissue}$ | No RT (N=7,7,4) | 2199 ± 655 | 1332 ± 622† | 1138 ± 537† |
| | 2Gy (N=7,7,6) | 2082 ± 1528 | 4984 ± 1685*† | 1785 ± 738¶ |
| | 10Gy (N=6) | 2193 ± 737 | 6793 ± 1932*† | 4723 ± 1943*§ |
| $WIS_{whole}$ | No RT (N=7,7,4) | 5050 ± 1653 | 3599 ± 2063 | 3007 ± 1411† |
| | 2Gy (N=7,7,6) | 4299 ± 2257 | 7469 ± 2426* | 3662 ± 1994 |
| | 10Gy (N=6) | 4168 ± 909 | 10195 ± 3604*† | 7041 ± 2392* |
| $PH_{tissue}$ | No RT (N=7,7,4) | 954 ± 259 | 571 ± 200† | 549 ± 260 |
| | 2Gy (N=7,7,6) | 902 ± 479 | 1817 ± 470** | 721 ± 257¶ |
| | 10Gy (N=6) | 909 ± 257 | 2090 ± 588*† | 1703 ± 650*§† |
| $PH_{whole}$ | No RT (N=7,7,4) | 1111 ± 261 | 745 ± 302† | 681 ± 322 |
| | 2Gy (N=7,7,6) | 1084 ± 489 | 2004 ± 474** | 871 ± 319¶ |
| | 10Gy (N=6) | 1043 ± 279 | 2282 ± 633*† | 1847 ± 630*§† |
| $PH_{blood}$ | No RT (N=7,7,4) | 2526 ± 367 | 2068 ± 949 | 1702 ± 856 |
| | 2Gy (N=7,7,6) | 2374 ± 639 | 2581 ± 645 | 1857 ± 590 |
| | 10Gy (N=6) | 2114 ± 497 | 2600 ± 841 | 2090 ± 599 |
| $v_{ec}$ | No RT (N=6,6,4) | 0.366 ± 0.124 | 0.321 ± 0.051 | 0.367 ± 0.06 |
| | 2Gy (N=7,7,6) | 0.344 ± 0.118 | 0.449 ± 0.045* | 0.359 ± 0.072¶ |
| | 10Gy (N=6) | 0.403 ± 0.091 | 0.509 ± 0.088* | 0.533 ± 0.109*§† |
| $K_{trans}$ | No RT (N=6,6,4) | 0.448 ± 0.129 | 0.328 ± 0.155 | 0.308 ± 0.13 |
| | 2Gy (N=7,7,6) | 0.543 ± 0.295 | 1.038 ± 0.302** | 0.496 ± 0.167¶ |
| | 10Gy (N=6) | 0.614 ± 0.22 | 1.746 ± 0.388**§† | 1.336 ± 0.214**§§† |
| $K_{ep}$ | No RT (N=6,6,4) | 1.27 ± 0.27 | 1.05 ± 0.52 | 0.82 ± 0.27 |
| | 2Gy (N=7,7,6) | 1.56 ± 0.56 | 2.31 ± 0.63* | 1.37 ± 0.4 |
| | 10Gy (N=6) | 1.55 ± 0.49 | 3.54 ± 1.09*† | 2.62 ± 0.77*§ |
| Vessel Density ($mm^{-2}$) | No RT (N=6,6,4) | 400 ± 42 | 380 ± 30 | 381 ± 43 |
| | 2Gy (N=6) | 409 ± 47 | 380 ± 39 | 365 ± 57 |
| | 10Gy (N=6) | 432 ± 30 | 337 ± 53*† | 275 ± 36*§†† |
| Mean Vessel Diameter (µm) | No RT (N=6,6,4) | 10.3 ± 1 | 10.5 ± 1.6 | 11.3 ± 2.6 |
| | 2Gy (N=6) | 10.7 ± 1.3 | 14.7 ± 1.5*† | 13 ± 2.7 |
| | 10Gy (N=6) | 9.4 ± 1.9 | 16 ± 1.9**†† | 20.2 ± 2.5*§††¶ |
| Single-vessel Blood flow (µm/s) | No RT (N=6,6,3) | 1299 ± 257 | 1259 ± 377 | 895 ± 459 |
| | 2Gy (N=6) | 1231 ± 70 | 2312 ± 405*† | 1852 ± 584 |
| | 10Gy (N=6) | 1420 ± 281 | 2096 ± 295*† | 2503 ± 536*†† |

[#]-PB samples were not taken in leukemic mice after RT as an appropriate number of cells could not be obtained from the maximum sampled blood volume

[†]-Significantly different from pretreatment time-point P<0.05

[¶]-Significantly different from 2 days post-treatment time-point P<0.05

* -Significantly different from no RT group P<0.05

[§]-significantly different from 2Gy treatment group P<0.05

[††]-Significantly different from pretreatment time-point P<0.001

[¶¶]-Significantly different from 2 days post-treatment time-point P<0.001

**-Significantly different from no RT group P<0.001

[§§-]Significantly different from 2Gy treatment group P<0.001

#### Table E2

| ALL Low Disease Burden | | Pretreatment | Day 2 Post-treatment | Day 5 Post-treatment |
|---|---|---|---|---|
| | | Mean ± Standard Deviation | Mean ± Standard Deviation | Mean ± Standard Deviation |
| % ALL in PB | No RT (N=9) | 5.5 ± 3.3 | 24.0 ± 9.8†† | 30.0 ± 8.4†† |
| | 2Gy (N=7) | 7.6 ± 4.3 | # | # |
| | 10Gy (N=7) | 6.6 ± 3.5 | # | # |
| $WIS_{tissue}$ | No RT (N=7) | 3347 ± 1986 | 1860 ± 895 | 1928 ± 1280 |
| | 2Gy (N=6) | 2797 ± 843 | 6706 ± 1469**†† | 2244 ± 1528¶¶ |
| | 10Gy (N=6) | 2825 ± 1042 | 7956 ± 1285**† | 6472 ± 2465*§† |
| $WIS_{whole}$ | No RT (N=7) | 6960 ± 3423 | 3820 ± 2271 | 3057 ± 2476† |
| | 2Gy (N=6) | 5849 ± 2067 | 11518 ± 3019*†† | 3729 ± 2752¶¶ |
| | 10Gy (N=6) | 4847 ± 1115 | 12224 ± 2105**† | 10047 ± 3358*§† |
| $PH_{tissue}$ | No RT (N=7) | 955 ± 396 | 848 ± 381 | 883 ± 505 |
| | 2Gy (N=6) | 846 ± 407 | 1577 ± 261*†† | 783 ± 342¶ |
| | 10Gy (N=6) | 941 ± 452 | 2806 ± 527**§†† | 2299 ± 522*§§† |
| $PH_{whole}$ | No RT (N=7) | 1339 ± 542 | 1072 ± 486 | 1011 ± 579 |
| | 2Gy (N=6) | 1245 ± 511 | 2188 ± 422*†† | 937 ± 437¶¶ |
| | 10Gy (N=6) | 1173 ± 430 | 3020 ± 406**§†† | 2548 ± 403**§§†† |
| $PH_{blood}$ | No RT (N=7) | 2400 ± 821 | 1805 ± 597 | 1530 ± 707† |
| | 2Gy (N=6) | 2427 ± 783 | 3061 ± 423* | 1716 ± 803¶ |
| | 10Gy (N=6) | 2123 ± 408 | 3267 ± 251**† | 2887 ± 525*§ |
| $v_{ec}$ | No RT (N=7) | 0.277 ± 0.081 | 0.289 ± 0.041 | 0.357 ± 0.066 |
| | 2Gy (N=6) | 0.229 ± 0.076 | 0.305 ± 0.024 | 0.304 ± 0.078† |
| | 10Gy (N=6) | 0.278 ± 0.062 | 0.513 ± 0.061**§§†† | 0.491 ± 0.083*§† |
| $K_{trans}$ | No RT (N=7) | 0.93 ± 0.247 | 0.685 ± 0.192 | 0.609 ± 0.222 |
| | 2Gy (N=6) | 0.833 ± 0.192 | 1.983 ± 0.433**† | 0.892 ± 0.363¶ |
| | 10Gy (N=6) | 0.902 ± 0.262 | 1.444 ± 0.149**† | 1.316 ± 0.399* |
| $K_{ep}$ | No RT (N=7) | 3.68 ± 1.51 | 2.37 ± 0.61 | 1.69 ± 0.5†¶ |
| | 2Gy (N=6) | 3.89 ± 1.16 | 6.6 ± 1.67*† | 3.04 ± 1.26¶ |
| | 10Gy (N=6) | 3.3 ± 0.82 | 2.85 ± 0.44§† | 2.65 ± 0.48* |
| Vessel Density ($mm^{-2}$) | No RT (N=6) | 419 ± 46 | 456 ± 40 | 529 ± 64†¶ |
| | 2Gy (N=6) | 461 ± 69 | 480 ± 70 | 481 ± 87 |
| | 10Gy (N=6) | 457 ± 72 | 356 ± 59*§† | 282 ± 63**§†¶ |
| Mean Vessel Diameter (μm) | No RT (N=6) | 12.3 ± 2.1 | 10.2 ± 1.2 | 10.2 ± 1.0† |
| | 2Gy (N=6) | 11.3 ± 1.5 | 13.3 ± 1.3*† | 9 ± 1.4†¶ |
| | 10Gy (N=6) | 11.7 ± 1.6 | 18.6 ± 1.9**§§† | 19.8 ± 4.2*§† |
| Single-vessel Blood flow (μm/s) | No RT (N=6) | 1292 ± 453 | 595 ± 302† | 494 ± 242† |
| | 2Gy (N=6) | 1340 ± 337 | 2312 ± 580**† | 1367 ± 596* |
| | 10Gy (N=6) | 1049 ± 352 | 2017 ± 668*† | 1204 ± 374* |

[#]-PB samples were not taken in mice bearing ALL after RT as an appropriate number of cells could not be obtained from the maximum sampled blood volume
[†]-Significantly different from pretreatment time-point P<0.05
[¶]-Significantly different from 2 days post-treatment time-point P<0.05
* -Significantly different from no RT group P<0.05
[§]-significantly different from 2Gy treatment group P<0.05
[††]-Significantly different from pretreatment time-point P<0.001
[¶¶]-Significantly different from 2 days post-treatment time-point P<0.001
**-Significantly different from no RT group P<0.001
[§§]-Significantly different from 2Gy treatment group P<0.001

**Table E3**

| ALL High Disease Burden (N=5) | Pretreatment | 2 days post 10Gy TBI | Paired t-test |
|---|---|---|---|
| | Mean ± Standard Deviation | Mean ± Standard Deviation | P value |
| % ALL in PB | 41.4 ±9.2 | # | # |
| $WIS_{tissue}$ | 1546 ± 192 | 5056 ± 1626 | **0.0091** |
| $WIS_{whole}$ | 2661 ± 363 | 7202 ± 2078 | **0.0054** |
| $PH_{tissue}$ | 872 ± 263 | 2295 ± 670 | **0.0041** |
| $PH_{whole}$ | 1033 ± 332 | 2398 ± 626 | **0.0028** |
| $PH_{blood}$ | 1365 ± 387 | 2491 ± 602 | **0.0029** |
| $v_{ec}$ | 0.387 ± 0.017 | 0.550 ± 0.091 | **0.0245** |
| $K_{trans}$ | 0.759 ± 0.315 | 1.241 ± 0.365 | 0.0698 |
| $K_{ep}$ | 1.95 ± 0.77 | 2.25 ± 0.51 | 0.3627 |
| Single-vessel Blood flow (μm/s) | 1045 ± 74 | 2217 ± 361 | **0.0010** |

[#-]PB samples were not taken in mice bearing ALL after RT as an appropriate number of cells could not be obtained from the maximum sampled blood volume

#### Table E4

| WT | | Pretreatment<br>Mean ± Standard Deviation | Day 2 Post-treatment<br>Mean ± Standard Deviation | Day 5 Post-treatment<br>Mean ± Standard Deviation |
|---|---|---|---|---|
| $WIS_{tissue}$ | No RT (N=7) | 3233 ± 1881 | 2887 ± 954 | 2227 ± 970 |
| | 2Gy (N=6) | 2593 ± 898 | 3612 ± 1504† | 1749 ± 462†¶ |
| | 10Gy (N=6) | 3496 ± 1649 | 6317 ± 1927* | 4938 ± 3232 |
| $WIS_{whole}$ | No RT (N=7) | 7982 ± 4007 | 7241 ± 3590 | 5603 ± 3411 |
| | 2Gy (N=6) | 6793 ± 2977 | 7492 ± 4395 | 4480 ± 2933 |
| | 10Gy (N=6) | 8935 ± 4124 | 10513 ± 3891 | 8392 ± 3770 |
| $PH_{tissue}$ | No RT (N=7) | 1097 ± 737 | 1043 ± 537 | 737 ± 399 |
| | 2Gy (N=6) | 697 ± 223 | 826 ± 340 | 569 ± 255¶ |
| | 10Gy (N=6) | 909 ± 257 | 2090 ± 588*§† | 1703 ± 650*§† |
| $PH_{whole}$ | No RT (N=7) | 1503 ± 772 | 1407 ± 671 | 1048 ± 583 |
| | 2Gy (N=6) | 1213 ± 497 | 1369 ± 736 | 891 ± 459† |
| | 10Gy (N=6) | 1609 ± 691 | 1929 ± 705 | 1610 ± 757 |
| $PH_{blood}$ | No RT (N=7) | 2669 ± 782 | 2367 ± 907 | 2072 ± 1245 |
| | 2Gy (N=6) | 2178 ± 626 | 2011 ± 854 | 1657 ± 968 |
| | 10Gy (N=6) | 2663 ± 834 | 2460 ± 629 | 1997 ± 647 |
| $v_{ec}$ | No RT (N=7) | 0.295 ± 0.096 | 0.337 ± 0.055 | 0.329 ± 0.074 |
| | 2Gy (N=6) | 0.254 ± 0.054 | 0.308 ± 0.033 | 0.307 ± 0.086 |
| | 10Gy (N=6) | 0.274 ± 0.031 | 0.422 ± 0.039*§§† | 0.392 ± 0.097 |
| $K_{trans}$ | No RT (N=7) | 0.725 ± 0.266 | 0.72 ± 0.21 | 0.793 ± 0.314 |
| | 2Gy (N=6) | 0.726 ± 0.126 | 1.249 ± 0.35*† | 0.688 ± 0.317¶ |
| | 10Gy (N=6) | 0.93 ± 0.348 | 2.062 ± 0.409**§† | 2.112 ± 1.334 |
| $K_{ep}$ | No RT (N=7) | 2.49 ± 0.57 | 2.18 ± 0.73 | 2.52 ± 1.05 |
| | 2Gy (N=6) | 2.92 ± 0.5 | 4.04 ± 1.03*† | 2.37 ± 1.28¶ |
| | 10Gy (N=6) | 3.45 ± 1.28 | 4.89 ± 0.9** | 5.26 ± 2.32 |
| Vessel Density ($mm^{-2}$) | No RT (N=6) | 291 ± 73 | 288 ± 41 | 273 ± 43 |
| | 2Gy (N=6) | 307 ± 41 | 300 ± 51 | 277 ± 40 |
| | 10Gy (N=6) | 290 ± 49 | 253 ± 26 | 200 ± 32*§†¶ |
| Mean Vessel Diameter (µm) | No RT (N=6) | 16.6 ± 1.9 | 15.6 ± 1.7 | 16.1 ± 1.9 |
| | 2Gy (N=6) | 16.7 ± 0.9 | 20.3 ± 2.2*† | 19.1 ± 3.2 |
| | 10Gy (N=6) | 16.2 ± 2.4 | 20.8 ± 1.9*† | 26.6 ± 4*§†¶ |
| Single-vessel Blood flow (µm/s) | No RT (N=6) | 2045 ± 485 | 1770 ± 352 | 1975 ± 445 |
| | 2Gy (N=6) | 1863 ± 371 | 2300 ± 521 | 2338 ± 628 |
| | 10Gy (N=6) | 1902 ± 550 | 2062 ± 600 | 1908 ± 455 |

†-Significantly different from pretreatment time-point P<0.05

¶-Significantly different from 2 days post-treatment time-point P<0.05
* -Significantly different from no RT group P<0.05
§-significantly different from 2Gy treatment group P<0.05
††-Significantly different from pretreatment time-point P<0.001
¶¶-Significantly different from 2 days post-treatment time-point P<0.001
**-Significantly different from no RT group P<0.001
§§-Significantly different from 2Gy treatment group P<0.001

### Table E5

| | No Disease | | ALL 0.5-20% PB | | ALL 25-55% PB | | AML 25-60% PB | | AML 60-90% PB | |
|---|---|---|---|---|---|---|---|---|---|---|
| | Mean ± Standard Deviation | (N) | Mean ± Standard Deviation | P value (N) | Mean ± Standard Deviation | P value (N) | Mean ± Standard Deviation | P value (N) | Mean ± Standard Deviation | P value (N) |
| $WIS_{tissue}$ | 3114 ± 1518 | (19) | 2940 ± 899 | 0.6835 (14) | 1380 ± 363.9 | **0.0001** (10) | 2280 ± 1279 | 0.1326 (10) | 1268 ± 673.5 | **0.0001** (9) |
| $WIS_{whole}$ | 7907 ± 3648 | (19) | 5525 ± 1621 | **0.0178** (14) | 2153 ± 671.8 | **<0.0001** (10) | 4417 ± 1776 | **0.0018** (10) | 3024 ± 1549 | **<0.0001** (9) |
| $PH_{tissue}$ | 955 ± 524 | (19) | 989 ± 468 | 0.844 (14) | 741 ± 230 | 0.1404 (10) | 926 ± 421 | 0.8726 (10) | 634 ± 286 | **0.04688** (9) |
| $PH_{whole}$ | 1445 ± 654 | (19) | 1293 ± 487 | 0.449 (14) | 865 ± 292 | **0.0028** (10) | 1099 ± 430 | 0.0999 (10) | 753 ± 328 | **0.001** (9) |
| $PH_{blood}$ | 2512 ± 749 | (19) | 2286 ± 496 | 0.3063 (14) | 1268 ± 356 | **<0.0001** (10) | 2256 ± 486 | 0.2779 (10) | 1809 ± 866 | 0.0554 (9) |
| $v_{ec}$ | 0.276 ± 0.067 | (19) | 0.274 ± 0.084 | 0.9625 (14) | 0.364 ± 0.055 | **0.001** (10) | 0.365 ± 0.114 | **0.0412** (10) | 0.37 ± 0.069 | **0.006** (8) |
| $K_{trans}$ | 0.79 ± 0.267 | (19) | 0.866 ± 0.205 | 0.361 (14) | 0.634 ± 0.272 | 0.1557 (10) | 0.593 ± 0.285 | 0.0865 (10) | 0.372 ± 0.173 | **<0.0001** (8) |
| $K_{ep}$ | 2.93 ± 0.89 | (19) | 3.39 ± 1.07 | 0.1989 (14) | 1.73 ± 0.64 | **0.0003** (10) | 1.61 ± 0.53 | **<0.0001** (10) | 1 ± 0.46 | **<0.0001** (8) |
| Vessel Density ($mm^{-2}$) | 296 ± 53 | (18) | 476 ± 78 | **<0.0001** (14) | 505 ± 52 | **0.0011** (4) | 413 ± 39 | **<0.0001** (10) | 396 ± 55 | **0.0019** (7) |
| Mean Vessel Diameter (μm) | 16.5 ± 1.7 | (18) | 11.4 ± 1.5 | **<0.0001** (14) | 10.1 ± 1 | **<0.0001** (4) | 9.9 ± 1.7 | **<0.0001** (10) | 11.1 ± 1.9 | **<0.0001** (7) |
| Single-vessel Blood flow (μm/s) | 1938 ± 454 | (18) | 1043 ± 511 | **0.0125** (5) | 357 ± 99 | **<0.0001** (4) | 1320 ± 243 | **<0.0001** (11) | 1020 ± 334 | **<0.0001** (7) |

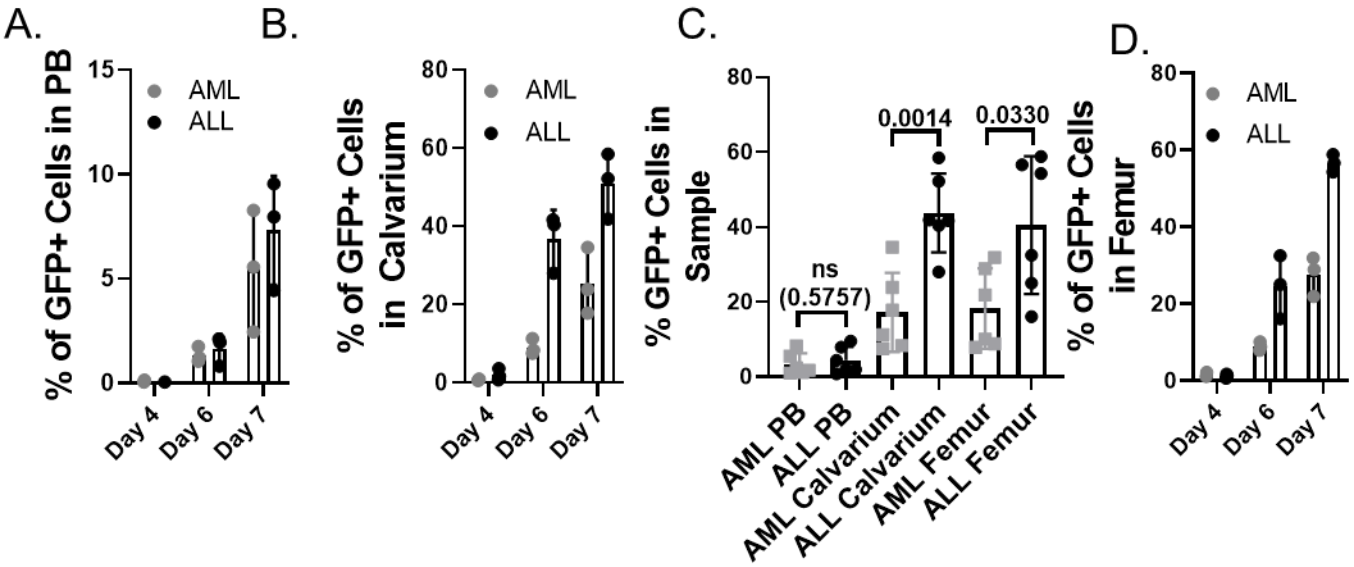


**Figure E1:**

The percentage of GFP+ ALL or AML cells in total cells for (A) peripheral blood, (B) crushed calvarium marrow, and (D) crushed femur marrow samples 4, 6, and 7 days post AML or ALL injection. (C) Comparison of the percentage of GFP+ ALL or AML cells in total cells for peripheral blood, crushed calvarium marrow, and crushed femur marrow samples taken from matching mice harvested 6 and 7 days after AML or ALL injection.

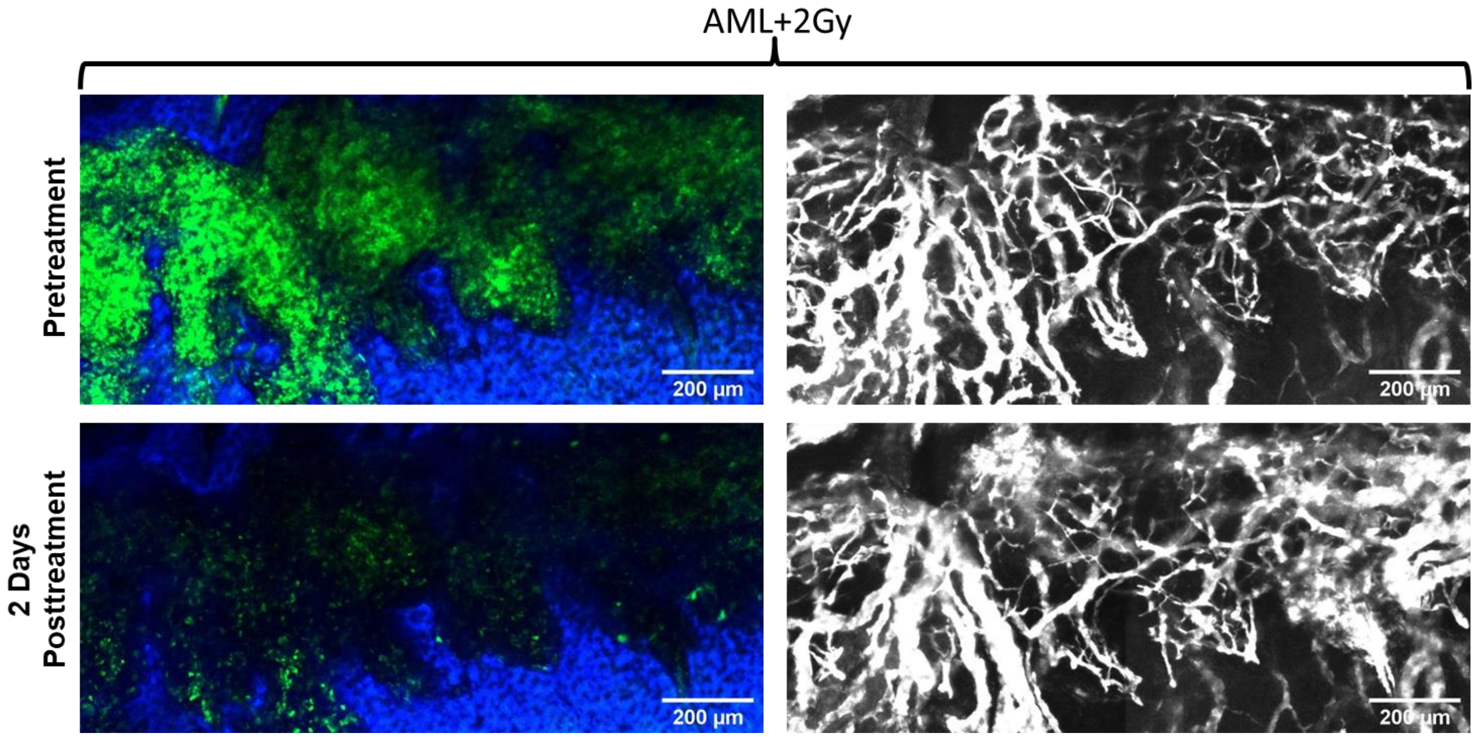


**Figure E2:**

Tiled, single-slice, co-localized images of GFP+ AML (green) and second harmonic generation of bone-collagen (blue) on the left. Maximum intensity projection images of Qtracker™ 655 labeled vascular blood pool (white) on the right. Images show the effect of 2Gy TBI on AML and the BMV.

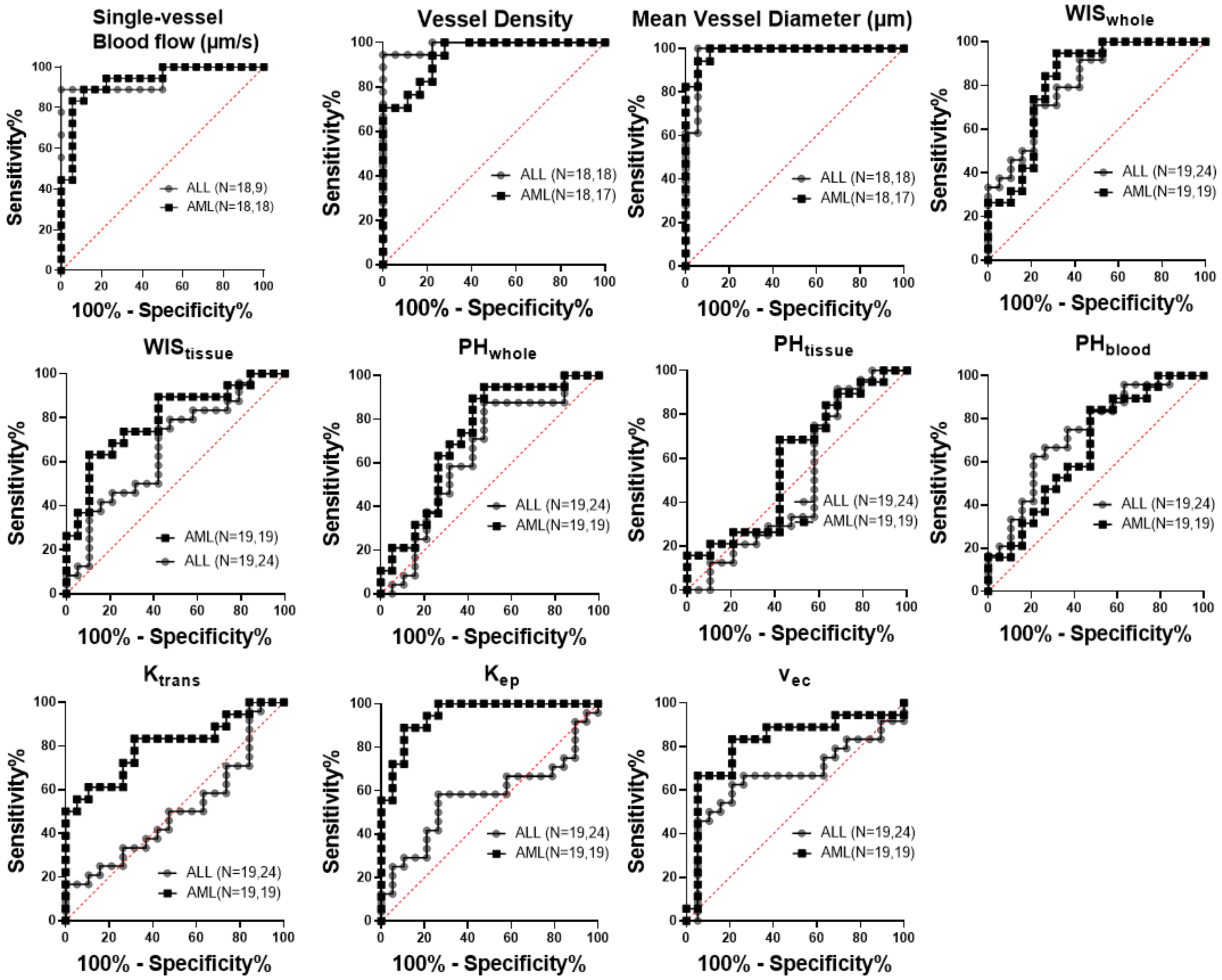


**Figure E3:**

ROC curves for L-QMPM imaging parameters for mice bearing AML and ALL imaged before treatment. Parameters include: Single-vessel blood flow, vessel density, mean vessel diameter, $WIS_{whole}$, $WIS_{tissue}$, $PH_{whole}$, $PH_{tissue}$, $PH_{blood}$, $K_{trans}$, $K_{ep}$, and $v_{ec}$.

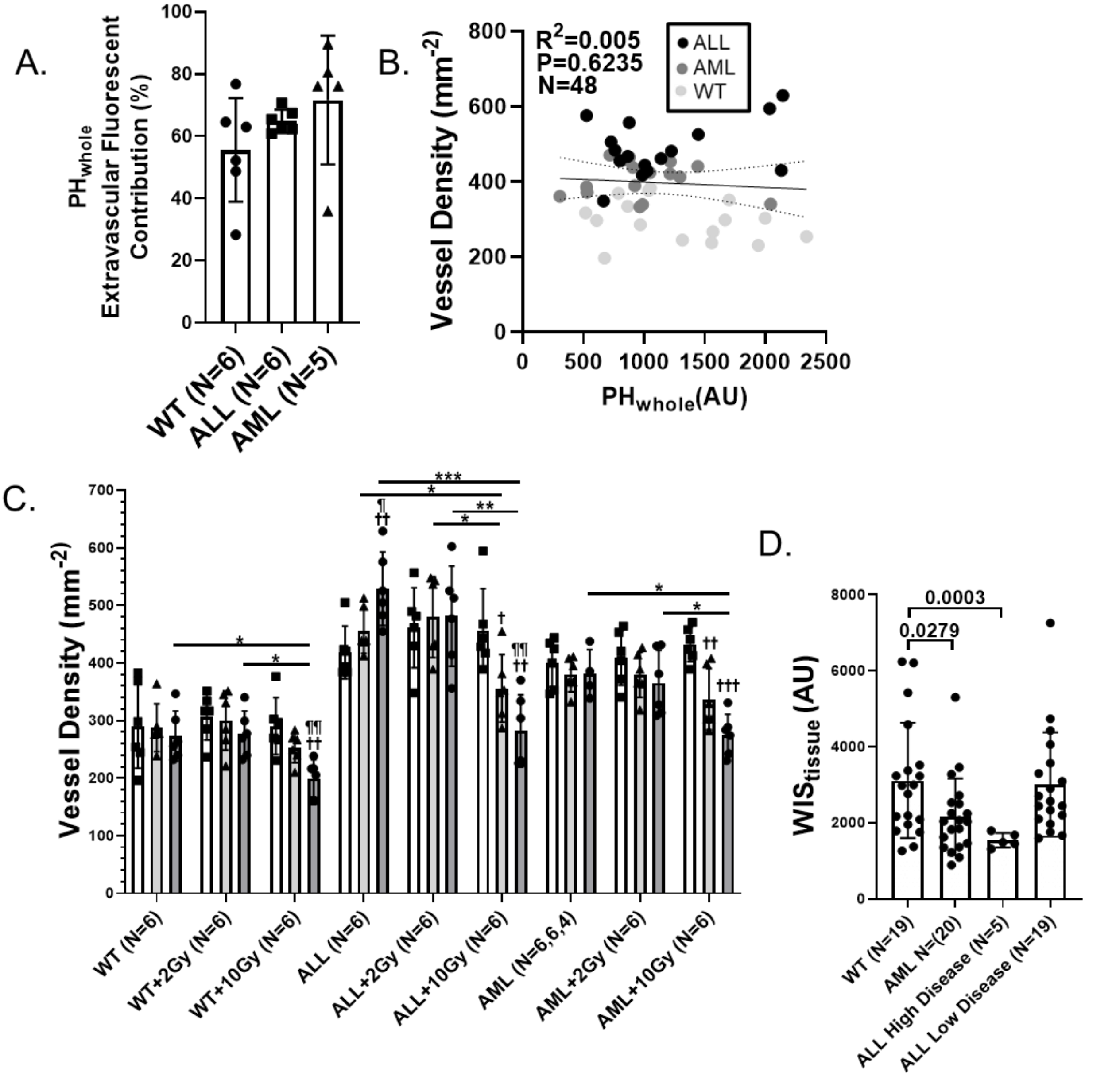


**Figure E4:**

(A) The percentage contribution of the extravascular tissue ROI region fluorescent signal to the total signal in $PH_{whole}$ for WT mice, and mice bearing AML or ALL. (B) A correlation plot of $PH_{whole}$ and vessel density showing no significant non-zero correlation between the parameters. (C) The effects of TBI on calvarium bone marrow vascular density for WT mice, and mice bearing ALL and AML (∗P < .05, ∗∗P < .01, ∗∗∗P < .001, ∗∗∗∗P < .0001, compared to pretreatment timepoint: †P < .05, ††P < .01, †††P < .001, ††††P < .0001, compared to 2 days posttreatment timepoint ¶P < .05, ¶¶P < .01, ¶¶¶P < .001, ¶¶¶¶P < .0001 Tukey's post hoc comparison). (D) The pretreatment values for $WIS_{tissue}$ in WT mice, mice bearing AML, mice bearing ALL at high disease burden, and mice bearing ALL at low disease burden from tables E4, E1, E3, and E2 respectively.
